# A historical specimen of False Lingzhi (*Ganoderma lucidum*) resolves a 245-year-old confusion within an important medicinal mushroom group

**DOI:** 10.64898/2026.05.13.724775

**Authors:** Torda Varga, Spike RJ Parker, Alessandro Agorini, Alexandra Dombrowski, Lewis Hadfield, A Martyn Ainsworth, David L. Hawksworth, Masoomeh Ghobad-Nejhad, Viktor Papp

**Affiliations:** Royal Botanic Gardens Kew, Richmond, TW9 3DS, UK; University of Gothenburg, Göteborg, 413 90, Sweden; The Natural History Museum, London, SW7 5BD, UK; Jilin Agricultural University, Changchun, 130118, Jilin Province, China; Geography and Environmental Science, University of Southampton, Southampton, SO7 1BJ, UK; Hungarian University of Agriculture and Life Sciences, Gödöllő, 2100, Hungary; Eötvös Loránd University, Budapest, 1117, Hungary

**Author notes:** These authors contributed equally (Torda Varga, Spike RJ Parker). Corresponding &.

**Keywords:** Traditional Chinese Medicine, DNA barcoding, Lingzhi, ITS, genomics

## Abstract

Plants and fungi are major sources of natural products beneficial to society, making the study of distinct species essential for discovering new drugs and bioactive compounds. The medicinal mushroom “Lingzhi” or “Reishi” (*Ganoderma lingzhi*) is widely used in traditional medicine and extensively studied for its bioactive triterpenoids, yet it is commonly identified as *Ganoderma lucidum*, the type species of the genus, which lacks a type specimen.
We sequenced a *G. lucidum* specimen preserved in the Kew fungarium, which agreed with the original description and was collected from wood of *Corylus avellana* in southern England. Using this reference specimen, we compiled genomic and ITS barcoding datasets to explore the genetic and geographic variation within this species.
We showed that *G. lingzhi* and *G. lucidum* diverged more than 12 million years ago and that all seven “*G*. *lucidum*” genomes deposited in public databases belong to other species. More than 1000 barcoding sequences showed that the widely used homology-based ITS barcoding is not working in this group, which can be mitigated by a phylogenetic placement approach. The 149 sequences assigned to *G. lucidum* with high confidence showed a Eurasian distribution and introductions to North and South America and Africa.
Our study underscores the importance of accurate species identification and provides guidance for a group of pharmaceutical and socially significant species. To further support future studies and the wider public in differentiating between *G. lingzhi* and *G. lucidum*, we propose using “False Lingzhi” as the English name for *G. lucidum*.

**Societal Impact Statement:** Traditional Chinese Medicine has expanded far beyond Asia, with growing markets in North America and Europe for supplements and functional foods. Lingzhi or Reishi (*Ganoderma lingzhi*), a well-known medicinal mushroom, is valued for its anti-inflammatory and anticancer properties. However, it is often misidentified with species that may not provide the same health benefits. This confusion poses risks to consumer safety, product regulation, and research. Here, we establish a reference using morphological and molecular tools for the most commonly misidentified species (*Ganoderma lucidum*) and propose the name “False Lingzhi” to distinguish it, supporting accurate identification, safer product development, and reliable research.

## Introduction

Accurate species delimitation provides a foundation for understanding the biology, ecology and potential of economically and epidemiologically important organisms. Species identity can have direct practical consequences, as illustrated by vector-specific malaria control strategies and crop improvement efforts such as coffee breeding, where trait variation is linked to morphologically similar but phylogenetically different species (Kweyamba et al., 2025; Davis et al., 2025). Species names are anchored by type specimens that serve as the objective reference for any application (Thomson et al., 2018). However, these types are not always represented by preserved biological material but instead by an illustration, limiting morphological and preventing molecular studies and, leading to uncertainties in application and causing confusion. In these cases, additional interpretative type specimens (e.g., epitypes) can be designated (Turland et al., 2025).

Medicinal mushrooms are attracting scientific and commercial interest due to their long history of use in Traditional Chinese Medicine (TCM) and the diverse bioactive properties of the secondary metabolites they produce (Barua et al., 2024). Among them, members of the genus *Ganoderma*, particularly *G. lingzhi* commonly known as “Lingzhi” or “Reishi” in East Asia, rank among the most valuable and extensively studied medicinal mushrooms (Bishop et al., 2015; Bhunjun et al., 2024; Karunarathna et al., 2025). *Ganoderma* species are a major focus of pharmacological and preclinical research investigating the bioactive effects of mainly two groups of compounds: triterpenoids and beta-glucans (Paterson, 2006; Du et al., 2024; Wu et al., 2024). For example, among many promising results (Klupp et al., 2016; Liang et al., 2019; Jafari et al., 2025), preclinical work has shown that *Ganoderma*-specific triterpenoids (ganoderic acid A and DM) applied in low concentration induced brain tumour cell death and reduced tumour volume in mice (Das et al., 2020).

Despite the pharmacological and economic interest in *Ganoderma* species, the type species of the genus, *Ganoderma lucidum* (Curtis) P. Karst. has only a graphic illustration as a type (lectotype) rather than a specimen (Curtis, 1781), leaving scientists without any reference biological material for centuries (Steyaert, 1961). The importance of the absence of biological reference material for this species is highlighted by the varying composition and concentration of triterpenoids across *Ganoderma* species and strains (Welti et al., 2015), alongside the confusion surrounding the common and Latin names of various medicinal *Ganoderma* species. The name *G. lucidum* has historically been widely applied to morphologically similar collections from different geographic regions (Papp et al., 2017) and phylogenetically distinct lineages have frequently been treated under this single name, while differences among taxa have often been overlooked (Moncalvo et al., 1995b; Moncalvo & Ryvarden, 1997). A prominent example of this taxonomic ambiguity is the concept of “lingzhi”, a culturally and economically important medicinal fungus that has long been treated as synonymous with *G. lucidum* (Papp et al., 2017; Loyd et al., 2018). However, phylogenetic studies have demonstrated that the East Asian concept of “lingzhi” does not correspond to the European *G. lucidum*, but instead represents a distinct and evolutionarily separate lineage within Ganoderma (Moncalvo et al., 1995a; Szedlay, 2002; Wang et al., 2009; Wang et al., 2012; Cao et al., 2012), a genus now resolved into ten major subclades (Sun et al., 2022). This separation is further supported by differences in chemical composition (Welti et al., 2015; Hennicke et al., 2016). Although European *G*. *lucidum* and the East Asian “lingzhi” lineage are now known to occupy separate phylogenetic subclades, the precise taxonomic circumscription of “lingzhi” remains a subject of debate (Yao et al., 2020; Yao et al., 2013; Zhou et al., 2015; Dai et al., 2017; Du et al., 2023), and the name *G*. *lucidum* is still frequently applied to this medicinal mushroom in both the scientific literature and commercial use. The historical taxonomic confusion surrounding these taxa (Papp, 2019; Papp, 2024; Dai et al., 2017) has contributed substantially to inconsistent species identification in the literature and sequence databases, with biological, pharmacological and genomic data often falsely attributed to *G*. *lucidum* (Papp et al., 2017; Loyd et al., 2018; Fryssouli et al., 2020).

Moreover, species boundaries within the *G*. *lucidum* complex remain insufficiently resolved (Cortina-Escribano et al., 2024). Resolving taxonomic uncertainties requires reliable diagnostic tools capable of distinguishing closely related lineages. The cryptic nature of fungi and their generally limited morphological differentiation have made molecular identification, often referred to as DNA barcoding, an essential approach (Kauserud, 2023). The most widely used barcoding region is the ITS region, which is easy to amplify and sequence, making it one of the most common DNA fragments in reference databases and the gold-standard barcoding target (Schoch et al., 2012). Yet, despite its known inconsistencies and the shift toward multi-gene methods (Taylor, 2011; Hibbett et al., 2016), the inherent limitations of ITS barcoding remain underemphasized (but see Bradshaw et al., 2023; Kauserud, 2023).

To address biological and taxonomic uncertainties surrounding medicinal *Ganoderma* species, we designate a well-preserved specimen of *G. lucidum* as an epitype that corresponds to the original description of *Boletus lucidus* (Curtis, 1781), the basionym of *G. lucidum*. We linked this enigmatic name to a biological specimen after 245 years, then generated new molecular data together with curated reference datasets, which allowed us to reassess species boundaries within the genus and evaluate the reliability of ITS-based barcoding frameworks. We further examined publicly available sequence and genomic resources associated with *G. lucidum* to clarify its taxonomic placement and explore the species’ biogeographic distribution. Finally, we discuss the implications of these findings for conservation, genetics, and pharmaceutical applications, in comparison with the often conflated East Asian “lingzhi” lineage.

## Methods

### Specimen selection and morphological study

The *Ganoderma* collection of the Royal Botanic Gardens, Kew (RBGK) fungarium was examined to identify a specimen suitable for the epitypification of *G. lucidum*. Selection criteria included southern British origin, occurrence on *Corylus* wood (consistent with the originally figured material), and morphological conformity with the original description. Macromorphological characters were documented from the preserved basidiome fragment. Pore density (pores per mm) was estimated from the pore surface using a stereomicroscope. Light microscopic observations were conducted using an Axio Imager A2 light microscope (Carl Zeiss, Germany). Sections were prepared by freehand sectioning, and at least three independent measurements were taken using AxioVision Release 4.8.2 software. Microscopic preparations were mounted in 5% KOH to rehydrate tissues and observe structural details, mounted in Melzer’s reagent (MR) or Congo red. For examination of crustohymeniderm cells, the lacquered crust layer was carefully removed with acetone prior to sectioning. Basidiospore measurements were based on 30 mature (i.e., released from basidia) spores per specimen. Basidiospore ornamentation was further examined using scanning electron microscopy (SEM). Spores were obtained by gently scraping the hymenial surface to release mature spores, which were then mounted onto aluminium stubs using double-sticky tabs. Stubs were sputter-coated with platinum (5 nm) using a Quorum Q150T ES sputter coater. Imaging was performed with a high-resolution, field-emission SEM (Hitachi Regulus 8230 High-end).

### Molecular work

Approximately 30 mg of hymenial tissue was sampled from the selected specimen. DNA was extracted following a PTB-based ancient DNA protocol (Latorre *et al*., 2020). Polymerase chain reaction (PCR) mixtures included 7.5 μL of RedTaq mix, 0.6 μL of 10 μM forward primer, 0.6 μL of 10 μM reverse primer, 1 μL of BSA, 4.3 μL of water, and 1 μL of template DNA. Six loci were targeted for amplification with the respective forward and reverse primers: ITS (ITS1-F/ITS4-R), ITS1 (ITS1-F/ITS2-R), ITS2 (ITS3-F/ITS4-R), and rpb2 (RPB2-OphF1/RPB2-OphR1). All ITS PCRs followed a standard 55.0°C thermal cycling protocol (White *et al*., 1990), whereas *rpb2* PCRs were run at an annealing temperature of 53.0 °C. 15 μL of PCR product was purified using the Macherey-Nagel NucleoSpin Gel and PCR Clean-up kit following the manufacturer’s instructions.

Purified PCR products were sequenced using the Sanger EZ workflow at GENEWIZ Europe. All sequences were trimmed with a 5% error probability limit, and consensus reads were generated by aligning higher quality (>50% HQ) forward and reverse reads.

### Compiling phylogenetic data

To place the *G. lucidum* epitype in a broader context, we compiled a “genus-level dataset” by including at least two specimens per subclade described by Sun *et al*. (2022), prioritising type specimens (Supplementary Table 1). The “genus-level dataset” included 35 specimens and six loci: full ITS (internal transcribed spacer) region (ITS1, 5.8S, ITS2), nuclear LSU (large subunit ribosomal RNA), nuclear SSU (small subunit ribosomal RNA), RPB2 (second-largest subunit of RNA polymerase II), TEF1 (translation elongation factor 1-alpha unit) and mitochondrial SSU.

To further disentangle the relationship among closely related species, we also compiled a “lucidum subclade dataset”, including 28 specimens from 10 species and seven loci: the loci included in the genus-level dataset and the RPB1 locus (the largest subunit of RNA Polymerase II).

### Maximum likelihood tree inference

Alignments of individual loci were generated with Muscle v.5.2 (Edgar, 2022) and manually curated with AliView v.1.30. software (Larsson, 2014). The boundaries of ITS1, 5.8S and ITS2 loci were determined on the aligned full-length ITS locus using the ITSx v.1.1.3 (Bengtsson-Palme et al., 2013) program. Aligned and edited sequences were trimmed to retain only parsimony-informative and constant sites and to remove gappy sites using a dynamically determined threshold in the clipkit v. 2.6.1 (Steenwyk et al., 2020) program. The supermatrix and partition file were generated from the individual loci using AMAS (Borowiec, 2016). Maximum likelihood (ML) tree inference was performed by using IQ-TREE v. 3.0.1 (Wong et al., 2026) by finding the best model using ModelFinder (Kalyaanamoorthy et al., 2017) and 100 standard nonparametric bootstrap replicates. For the “lucidum subclade dataset”, the tree search parameters were tuned to be more thorough by setting the number of initial parsimony trees to 250 and the top 80 parsimony trees were optimised with ML nearest neighbour interchange (NNI) search, considering all NNIs to initialise a 20-candidate set for the ML search. The log-likelihood epsilon was set to 0.00001 for higher precision. Statistical species delimitation was performed on this tree using the mPTP webserver v. 0.2.5 (Kapli et al., 2017) using the *Ganoderma shanxiense* clade as an outgroup. A Bayesian analysis was performed by using an ML estimation starting point and generating 500,000 MCMC steps with 10% burn-in and sampling every 5000.

### Molecular dating

To infer the divergence times of the main *Ganoderma* clades, we transformed the genus-level tree branch lengths into time units using the treePL v1.0 (Smith & O’Meara, 2012) phylogenetic penalised-likelihood program. We set the root of the tree to the minimum and maximum ages of 18 and 36 myrs based on the *Ganodermites lybicus* fossil (Fleischmann et al., 2007). Parameters were initialised using the prime option, followed by a leave-one-out cross-validation to determine the optimal smoothing parameter. The time-calibrated tree was inferred by using a smoothing parameter of 10 and the thorough option. Phylogenetic trees and chronograms were generated by using the Ggtree v. 3.16.3 (Xu et al., 2021), Deeptime v.2.3.1 (Gearty, 2025) and ape v.5.7-1 (Paradis & Schliep, 2019) R packages.

### ITS-based barcoding

To measure ITS sequence divergence, we calculated P-distance based on the global alignment of the full-length ITS locus (ITS1, 5.8S, ITS2), using the *dist.dna* function of the ape v. 5.7-1 R package (Paradis & Schliep, 2019) with the model = “raw” parameter. We calculated local percentage identity distances by performing all-versus-all local BLAST between the ITS sequences of the “lucidum subclade dataset” using the Blast v. 2.16.0 program (Altschul et al., 1990). In this analysis, to obtain a more realistic percentage identity value without bias from missing data, we imputed eight nucleotides in the ITS sequence (positions 37–44 in the 5.8S region) of the epitype that were conserved across the entire *Ganoderma* genus.

### ITS phylogeny

To assess whether different alignment methods and evolutionary models could help resolve the ITS tree, we performed 32 ML phylogenetic analyses. We generated Muscle- or PRANK-based alignments, with and without overhanging SSU and LSU sequences on the ITS. The Muscle alignment was prepared as mentioned above, while the PRANK alignment was generated with the +F argument using the Prank v.170427 program (Löytynoja, 2013). InDel coding was done by using the *code.simple.gaps* function of the Phyloch v. 1.5-5 R package (Heibl, 2008) after trimming the alignments with Clipkit as above. Maximum likelihood trees were inferred as above, but the perturbation strength was set to 0.2. We also examined whether thinning the alignment to half the species, keeping only the ingroup and the longest sequences, could improve phylogenetic inference. To avoid biases caused by alignment inaccuracies, we also jointly inferred the alignment and the tree in a Bayesian framework implemented in baliphy v.4.1 (Redelings, 2021) using the default rs07 InDel model (Redelings & Suchard, 2007). Finally, we inferred phylogenies that included *RPB2* in addition to ITS. We assessed phylogenetic inference by comparing median bootstrap values, Robinson-Foulds (RF) distances, and species monophyly with the species tree (i.e., multilocus phylogeny). RF distances were calculated by the *RF.dist* function of the phangorn v. 2.11.1 package (Schliep, 2011). To assess the performance of the ITS sequence alignment, we estimated a machine-learning-based difficulty value for phylogenetic inference using Pythia v.2.0.0 (Haag et al., 2022).

### Genome tree

Altogether, 24 genome assemblies were retrieved from the NCBI Genome database in May 2025, including 10 species within the genus *Ganoderma* and one outgroup taxon, *Sanguinoderma infundibulare* (Table S1). Orthologous protein sequences were extracted from the genome assemblies by using the BUSCO v5.8.3 (Tegenfeldt et al., 2025) with the OrthoDB v10 polyporales_odb10 lineage database. To keep only the single-copy genes present in all genomes, we used the BUSCOfilter script (Hill, 2024) with the “common” mode. The single-copy orthologs were aligned individually using MAFFT v.7.526 with the L-INS-i algorithm (Katoh & Standley, 2013). Poorly aligned or ambiguous regions were trimmed using trimAl v1.5.1 with “automated1” setting (Capella-Gutiérrez et al., 2009). Trimmed alignments were concatenated into a supermatrix using AMAS (Borowiec, 2016). Phylogenetic inference was conducted using IQ-TREE v3 (Wong et al., 2026) under the partition model MFP+MERGE (Kalyaanamoorthy et al., 2017). Node support was evaluated with 1000 ultrafast bootstrap replicates (Hoang et al., 2018).

We also placed the genome assemblies into the genus-level tree. To do this, we extracted seven barcoding sequences (ITS, LSU, nSSU, mtSSU, *RPB1*, *RPB2*, *TEF*) by using a BLAST-based tool (Hill, 2021). We used the *G. lucidum* Cui 14404 specimen as a reference for all except the *RPB1* locus, for which we used the sequence of the *G. carnosum* MUCL 49464 specimen. If a BLAST search returned multiple sequence fragments, we aligned them in AliView and, if necessary, assembled them to obtain the most comprehensive sequence possible. If multiple variants were presented, we prioritised the most common variants within the genome. The ML tree was inferred as described at the “lucidum sublcade dataset” tree inference.

### *Ganoderma lucidum* ITS database assessment

We created an ITS database that could contain all possible *Ganoderma lucidum* sequences using three approaches: (1) sequences related to GenBank taxon identifiers of the species, (2) BLAST searches of the GenBank nucleotide database, and (3) targeted searches for corresponding sequences in the UNITE database. *Ganoderma lucidum* ITS sequences were downloaded by the “1_Download_seqs.R” custom script (Hodgson & Varga, 2023) using the Biostrings v.2.62.0 (Pagès et al., 2021), rentrez 1.2.3 (Winter, 2017), restez v. 2.1.4.9000 (Bennett et al., 2018) and seqinr v.4.2-30 (Charif & Lobry, 2007) packages. We searched for sequences linked to the “1127707”, “1077286”, “1077064”, “939174” and “5315” taxonomic IDs and only sequences longer than 50 bp were downloaded. We performed a local BLAST search against a mirrored GenBank nucleotide database (created on the 25 May 2025) using BLAST v. 2.16.0 (Altschul et al., 1990), and a web-based search (performed on the 12th of February 2026) to ensure that all sequences were included. In both searches, a 96% identity threshold was applied to include a sufficiently broad set of sequences. Finally, we downloaded sequences associated with the UNITE SH0762718 species hypothesis (Abarenkov et al., 2024; Kõljalg et al., 2024). The sequences from the different approaches were combined and used for subsequent phylogenetic placement.

### Phylogenetic Placement and Geographic Distribution

Phylogenetic placement can be performed on an unpartitioned phylogenetic model. Thus, to test which unpartitioned model approximates the species tree best, we inferred an ITS tree with overhanging LSU and SSU sequences and four Gamma rate-heterogeneity models (2, 4, 6, and 8 rate categories). We aligned the ITS sequences downloaded above to the reference alignment used using mafft v7.526 (Katoh & Standley, 2013) with the --add and --keeplength arguments. Then, epa-ng v0.3.8 (Barbera et al., 2019) was used, with the best-fitting model selected by MFP IQ-TREE: HKY[1.0/6.2696]+FU[0.2125/0.2427/0.2516/0.2932]+IU[0.8086]+G4[0.4791]. We turned off pre-masking and heuristic pre-placement of sequences. We included placements with a minimum likelihood weight of 0.001 and limited the output to 100 sequences per placement. We transformed the resulted json placement file into a csv format using the to_csv command of guppy developmental program (Matsen et al., 2010). Because *G. lucidum* was not resolved as monophyletic in the reference ITS tree, we grouped branches into four categories: (1) non-*G. lucidum* (2) *G. lucidum* backbone (3) *G. lucidum* clade A, and (4) *G. lucidum* clade B. Then we calculated cumulative likelihood weight (cLw) for each category by summing the likelihood weights of corresponding branches and classified sequences by allowing some uncertainty as follows. We classified sequences as *G. lucidum* clade A or clade B if the cLw were higher than 0.5 of branches related to clade A or B, respectively and at the same time, if adding the *G. lucidum* backbone branches exceeded 0.8 cLw. If cLw were higher than 0.8 for the sum of the three categories, but none of the clades exceeded 0.5 cLw, we classified sequences as *G. lucidum* without clade specification (i.e., Clade AB). If the cLw were higher than 0.8 for non-*G*.*lucidum* branches, the sequences were classified as not *G. lucidum,* and if none of the above were true, sequences were classified as unidentifiable.

To generate a spatial distribution map, coordinates were manually georeferenced from the available metadata on NCBI using Google Maps (maps.google.com). For coordinates with only country-level information, a random coordinate within the country was chosen using a random coordinate generator (https://www.latlong.net/random-coordinate-generator). These points, except for random points from Russia, were plotted in R (version 4.5.3) using the ggplot2 v. 4.0.2 (Wickham, 2009) and rnaturalearthdata v. 1.0.0 packages (South et al. 2026).

## Results

### *Designating* an epitype for *Ganoderma lucidum* after 245 years

To resolve the 245 years of uncertainty surrounding the medicinal properties and taxonomy of *G. lucidum*, caused by the absence of biological material linked to the lectotype, we sought a specimen that matched the original description, including the geographic location, the host tree on which it grew, and its morphology (Fig. 1, Box 1).

**Figure 1.**
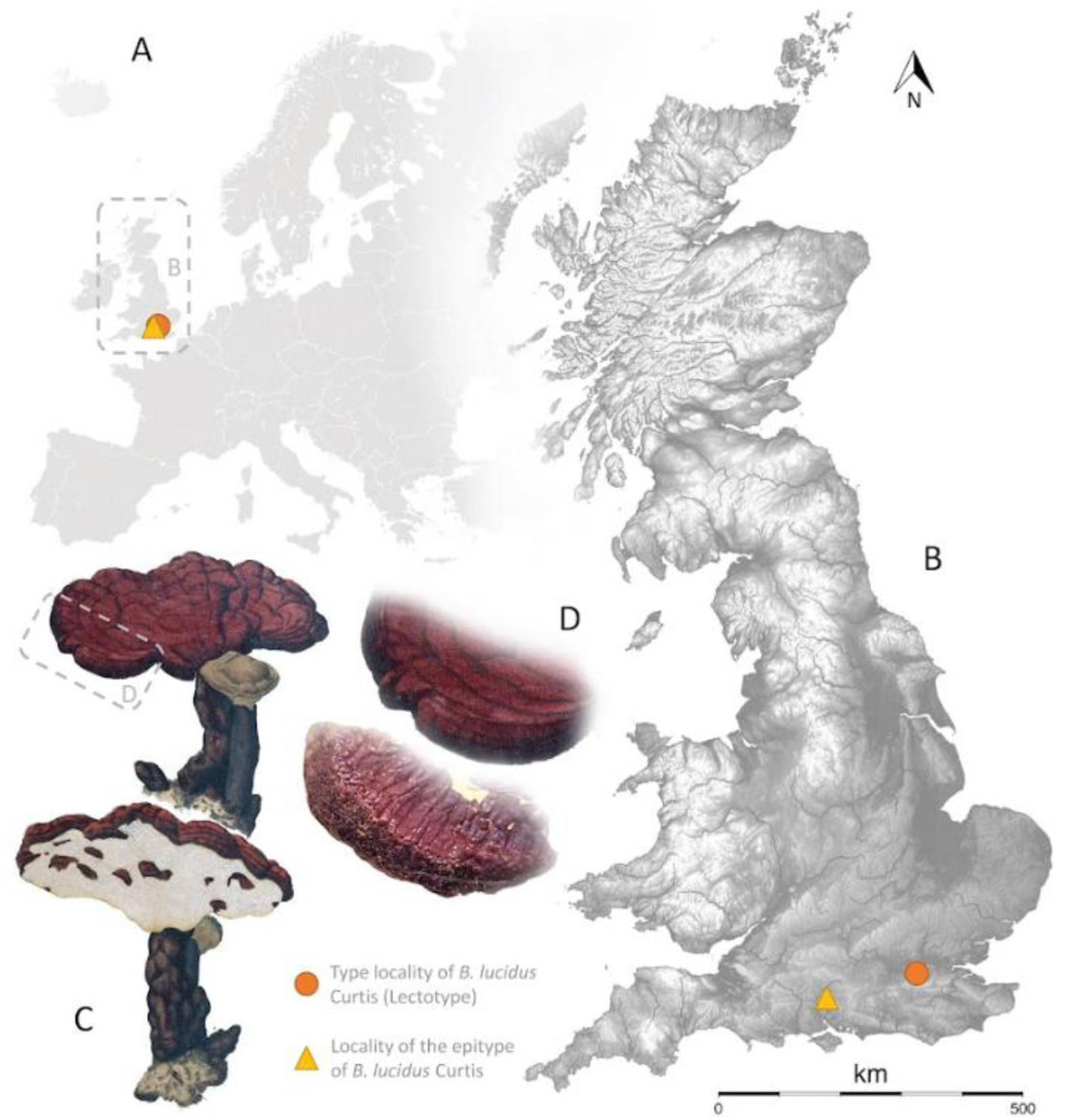
The macromorphology and locality of the lectotype and the newly designated epitype of *Ganoderma lucidum*. **A-B.** The locality of the lectotype (circle) and the epitype (triangle). **C.** The illustration of the original material. **D.** The macromorphology of the lectotype (top) and the epitype (bottom) is similar.

#### Box 1. The formal nomenclature of *G. lucidum* as interpreted here.

**Ganoderma lucidum** (Curtis) P. Karst., *Rev. Mycol. (Toulouse)*, **3** (9): 17 (1881). “False Lingzhi”

*Boletus lucidus* Curtis, *Fl. Londin.* **2** (4): 72^1^ (1781); nom. sanct. (Fr., *Syst. mycol*. **1**: 353; 1821, as *Polyporus lucidus*).

Type: Curtis, *Fl. Londin.* **2** (4): pl. 224 (1781) – lectotype (design. Steyaert in *Taxon* **10**: 251, 1961); United Kingdom, Hampshire, nr Winchester, Crab Wood, on wood [coppice stool] of *Corylus avellana*, 4 Apr. 2004, *P.J. Roberts,* K-M000121946 – epitype designated here, IF XXXXXXXX.

^1^This work is unpaginated, and the “72” is an index number allocated to it by Curtis; the text appears immediately before pl. 224; see Stevenson et al. (1961) for detailed bibliographic information on this work.

Attempts to recollect *G*. *lucidum* at the original type locality, Peckham Common, and at the adjacent Nunhead Cemetery on 8 October 2025 were unsuccessful, and no *Corylus*, the tree species associated with the lectotype, was observed. In addition, local fungal records indicated no documented occurrence of *G. lucidum* in the area since 1989 (personal communication by Liam Nash & Clifford Davy, 2025). A subsequent revision of the *Ganoderma* collections in the Kew (K-M) fungarium identified a single British specimen [K-M000121946] collected from *Corylus* wood in Crab Wood (Hampshire, England) (Figs. 1 and 2). Despite being labelled as *G. pfeifferi*, a species largely restricted to *Fagus*, it exhibited the diagnostic features traditionally associated with the European concept of *Ganoderma lucidum*, thus we designated this specimen as the epitype of *Boletus lucidus* Curtis, the basionym of *Ganoderma lucidum* (Curtis) P. Karst. (Box 1), here.

**Figure 2.**
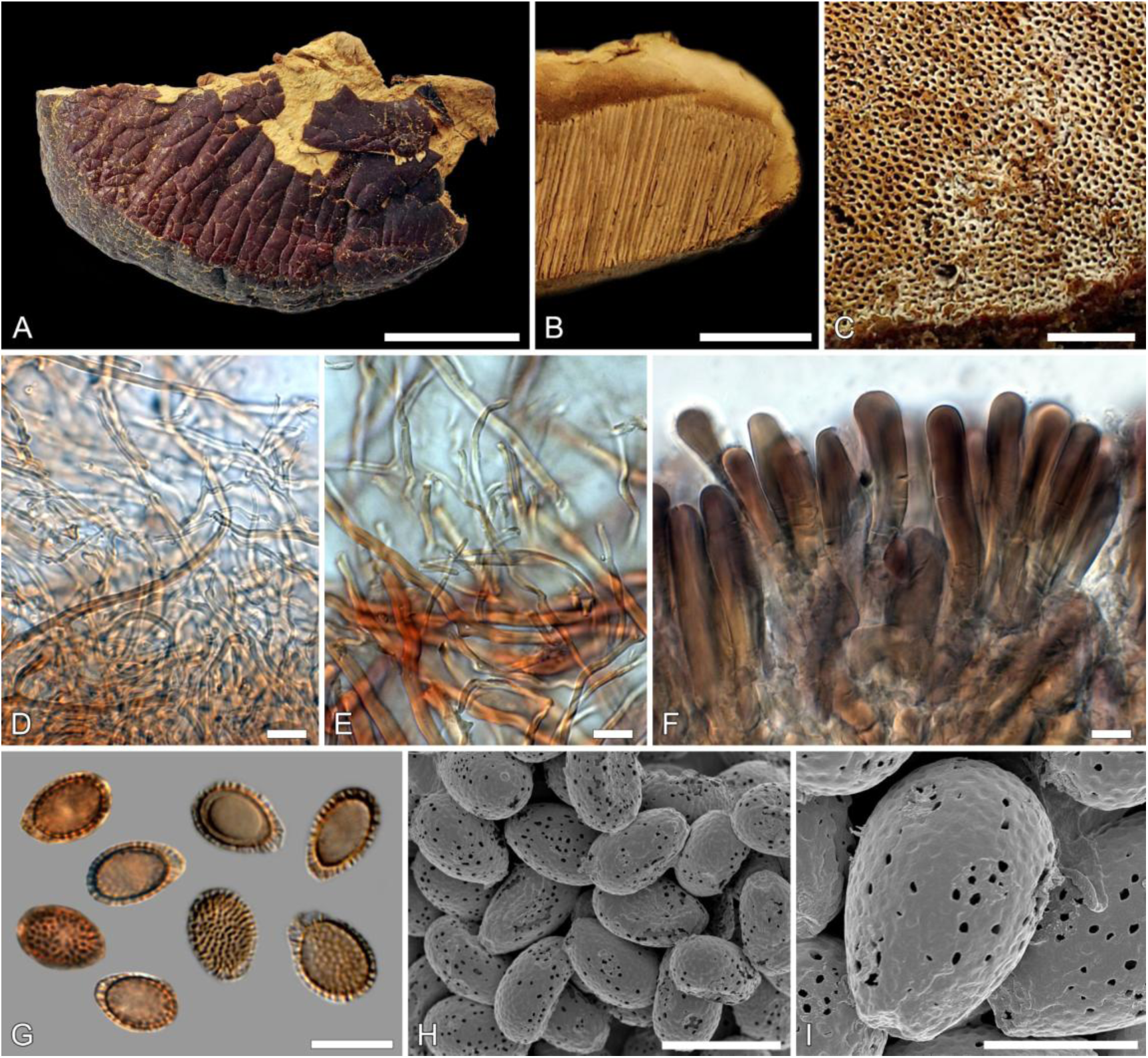
Macro- and micromorphological features of *Ganoderma lucidum* (epitype, K-M000121946). A) Upper surface of the basidiome; B) basidiome in cross section; C) pore surface; D) hyphal structure in the context; E) hyphal structure in the trama; F) pileipellis with crustohymeniderm cells; G) basidiospores under light microscopy; H, I) basidiospores under scanning electron microscopy (SEM). Scale bars: 1 cm (A); 5 mm (B); 1 mm (C); 10 µm (D–I).

The preserved fragment of an annual basidiome shows a laccate, reddish-brown pileal surface with an acute margin (Fig. 2A). The context is corky to woody and distinctly duplex, with a darker basal zone and a pale upper layer, the latter clearly distinguishing from *G. pfeifferi* the original label of the specimen (Fig. 2B). The pore surface is light brown, with pores circular to slightly angular, 4–5 per mm (average pore size is 170 µm) with thick dissepiments (Fig. 2C). The hyphal system is trimitic, with clamped generative hyphae, arboriform skeletal hyphae and binding hyphae typical of European laccate *Ganoderma* species in the *G*. *lucidum* complex (Fig. 2D,E). The pileipellis forms a well-developed crustohymeniderm composed of clavate to subcylindrical terminal cells with moderately thickened, pigmented walls arranged in a palisade-like layer (Fig. 2F). Basidiospores are ellipsoid, double-walled, with a truncate apex and ornamented endospore (Fig. 2G–I). Measurements based on 30 mature spores yielded dimensions of (10.5–)10.9–11.8(–11.9) × (6.8–)7.1–7.8(8.1) µm, average: 11.3 × 7.4 μm, Q= 1.4–1.6, Qav=1.5 μm (Table S2). SEM observations confirmed the characteristic verrucose endospore surface (Fig. 2H,I).

### The new biological material helps to disentangle molecular-based species boundaries

To further characterize the identity of the new type specimen, the single specimen which now fixes the application of the name, by using molecular tools, we sequenced the ITS (GenBank accession number: PZ166757) and *RPB2* (GenBank accession number: PZ179979) loci. We reconstructed the entire ITS region from two fragments, except for eight nucleotides (positions 37–44 in the 5.8S region) that are conserved at the family level. We inferred a genus-level ML tree using six ribosomal RNA and protein-coding genes, which clearly demonstrated that the designated *G. lucidum* epitype belongs to the “lucidum” subclade (subclade IX, BS = 100; Fig. 3/A). The monophyly of the ten subclades was highly supported (BS > 80) except in the case of subclade VIII (BS = 54). The relationships among the subclades were generally poorly resolved, with BS values ranging from 35% to 81% along the backbone of the genus. Only the clustering of subclades I-VI with subclade VI in a basal position received high support (BS = 81), whereas other relationships remained poorly supported. *Ganoderma resinaceum* and *G. linghzi* are also frequently misidentified as *G. lucidum*, but the genus-level tree reinforced these specimens’ position within the subclade VII (BS = 81). Based on the time calibration of the genus-level tree, we further showed that *G. lucidum* and *G. lingzhi* are evolutionarily highly distinct species, separated by at least 12 Myr.

**Figure 3.**
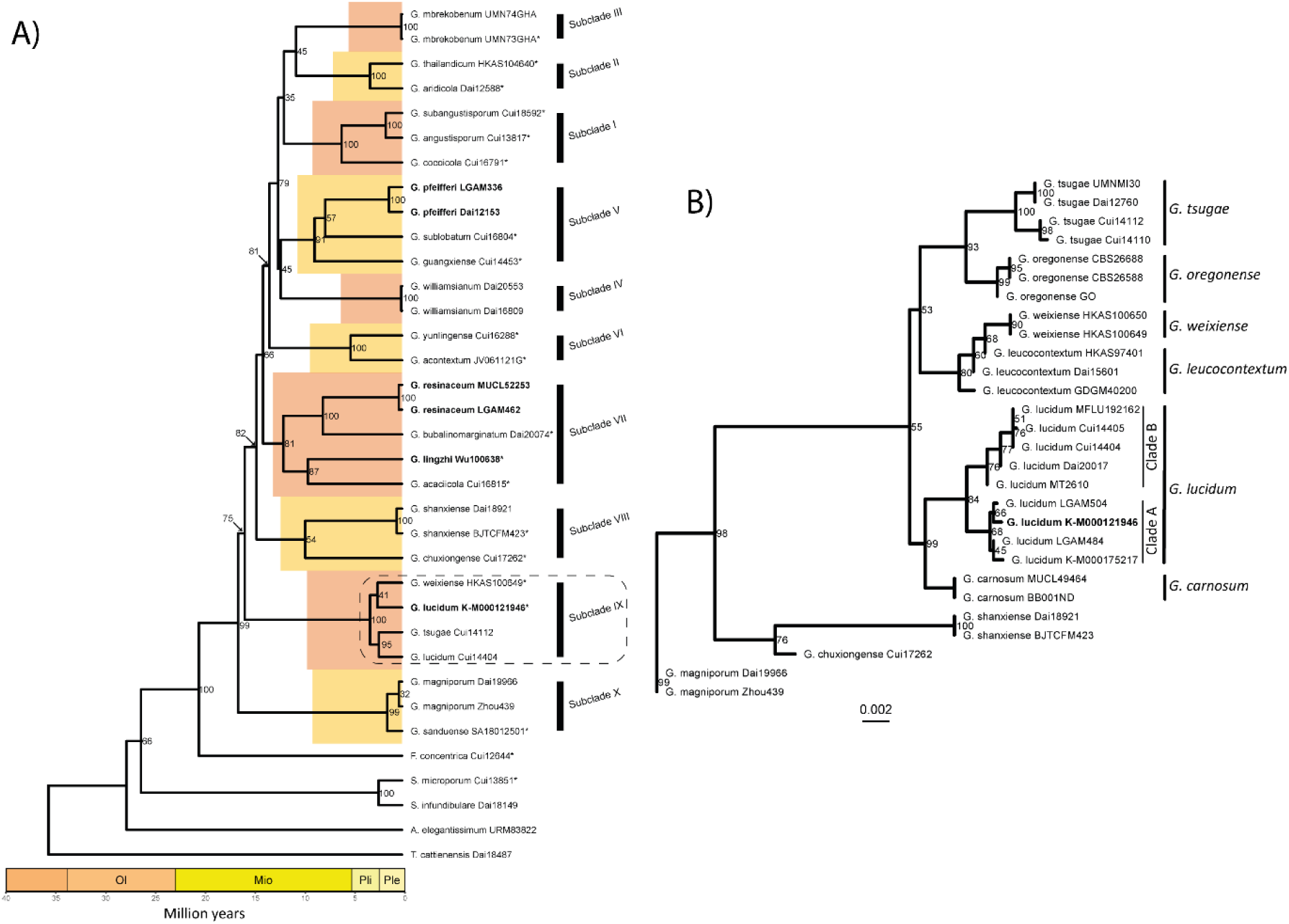
Phylogenetic position of *G. lucidum*. **A)** Genus-level chronogram based on six loci. Values at the nodes denote nonparametric bootstrap values. Type specimens were marked with an asterisk. Bolded names indicate that *G. pfeifferi* was the original label of the epitype K-M000121946, while *G. resinaceum* and *G. lingzhi* are commonly conflated with *G. lucidum*. **B)** The higher resolution phylogeny of the “lucidum” subclade IX based on seven loci. The *G. lucidum* epitype specimen is highlighted in bold.

To investigate the position of the epitype specimen at higher resolution, we inferred an ML tree using seven loci and including multiple specimens from all accepted species within subclade IX (Fig. 3/B). The monophyly of the species was well supported (BS > 84%), except for *G. leucocontextum*, which was paraphyletic within a clade that included *G. weixiense* specimens (BS = 80). The monophyly of *G. lucidum* specimens is supported (BS = 84%), while the sister species relationship of *G. lucidum* and *G. carnosum* is well resolved (BS = 99%). Two moderately well-supported subclades were further resolved within *G. lucidum*, one containing the new epitype (Clade A, BS = 68%) and the other containing Asian-origin specimens (Clade B, BS = 76%). Finally, despite strong phylogenetic and morphological support for species boundaries, modelling species delineation with mPTP did not detect multiple species within clade IX (Fig. S1).

### Evolutionary modelling is essential for accurate ITS barcoding

Because ITS barcoding is a widely used tool for species delineation in fungi, we analysed ITS sequence divergence and variation within the “lucidum subclade”, with particular emphasis on *G. lucidum*. We did not find an ITS sequence-similarity threshold that clearly distinguishes *G. lucidum* from other species. Yet, local alignment-based similarities (i.e., BLAST) were more sensitive to species boundaries (Fig. 4/A), whereas the global alignment-based p-distance showed lower resolution (Fig. S2). We determined that 98.86% identity is the lowest across the *G. lucidum* sequences using BLAST, whereas it is 97.0% using p-distance. However, both statistics showed a high degree of similarity between *G. lucidum* specimens and other species. For example, the MT-2610 specimen showed exceptionally high BLAST percentage identities to specimens of *G. tsugae* (100%), *G. oregonense* (99.5%), *G. weixiense* (99.1%), *G. leucocontextum* (99.1%) and *G. carnosum* (98.9%).

**Figure 4.**
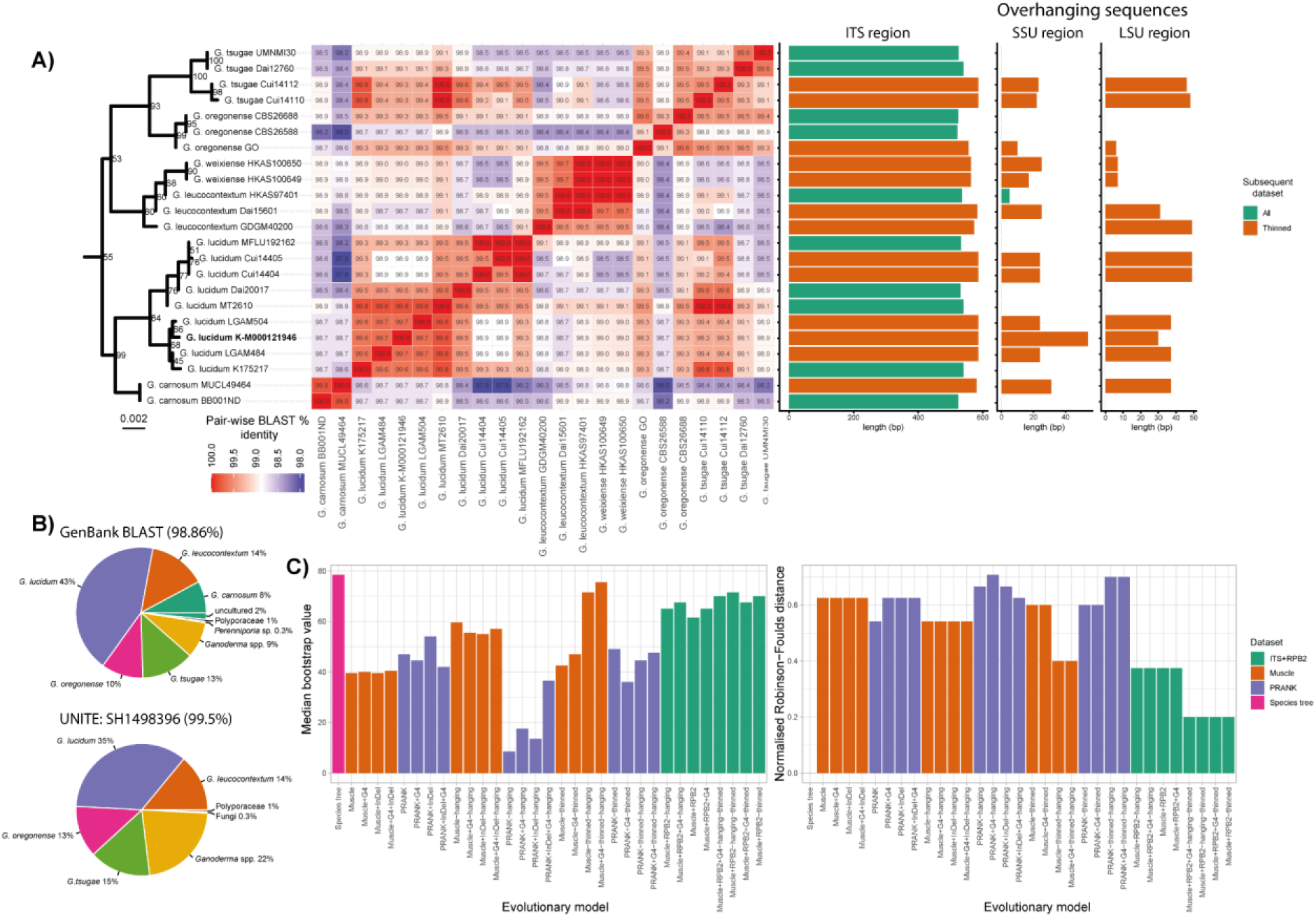
Assessing ITS barcoding of species within the “lucidum” clade with a special focus on *G. lucidum*. **A)** Sequence similarity based comparison of ITS sequences within the “lucidum” clade. Heatmap shows all-versus-all BLAST % identity comparisons. Barcharts depict the length of each ITS sequence and the overhanging SSU and LSU partial sequences. Colour codes of the bar charts indicate a subgroup of sequences (“thinned” dataset) that were used in some subsequent evolutionary modelling in panel C. **B)** The results of sequence similarity searches based on the *G. lucidum* epitype ITS in the GenBank (top) and UNITE (bottom) databases. **C)** Assessing different datasets and evolutionary models for inferring an ITS tree in comparison to the species tree. Muscle and PRANK denote two different multi-sequence alignment approaches of ITS. InDel indicates the modelling of insertion/deletion patterns. Thinned datasets represent sequences labelled as “thinned” in panel A. G4 denotes a Gamma model with four discrete categories.

Performing a BLAST search in the NCBI GenBank database using the 98.86% identity threshold returned 291 sequences, but 149 were labelled as a different species (Fig. 4/B). The UNITE database, which contains sequence-similarity-based clusterings of ITS sequences that provide species hypotheses, was also inconclusive regarding *G. lucidum*. The different similarity thresholds we applied yielded 364 sequences (SH1498396, 0.5%), 393 sequences (SH1088728, 1%), and 401 sequences (SH0762718, 1.5%). 75% of the sequences under the most stringent species hypothesis (SH1498396) were labelled as different species, including *G. tsugae* (15%), *G. leucocontextum* (14%), and *G. oregonense* (13%) (Fig. 4/B).

Next, we assessed whether phylogenetic modelling could help with ITS-based species delineation by seeking a model and a multisequence alignment that yielded the highest similarity to the species tree (i.e., multi-gene phylogeny), had the highest median bootstrap value, and resolved monophyletic species. Out of the 24 ML ITS-based phylogenies, the tree where half of the specimens were removed (i.e., thinned alignment), SSU and LSU overhanging fragments were kept, and the rate variation across sites was modelled, performed the best, with a median bootstrap value of 75.5%, normalised RF-dist of 0.4 compared to 80% bootstrap value of the species tree. If we considered phylogenies without thinning, the Muscle-based alignment with SSU and LSU overhanging fragments proved to be the best (median BS = 59.5, normalised RF = 0.54). Interestingly, modelling InDel evolution did not improve phylogenies in most cases, probably due to the low number of such characters (four in Muscle and three in PRANK alignments), and the InDels were informative only in relation to the outgroup. Nevertheless, neither the full nor the thinned alignment-based inferences could resolve the monophyly of *G. lucidum*, because two *G. tsugae* specimens (Vouchers: Cui 14112, Cui 14110) clustered together with the *G. lucidum* specimens. This could not be resolved by jointly inferring the alignment and the phylogeny in a Bayesian framework (Fig. S3). However, including the *RPB2* protein-coding gene fully resolved both *G. lucidum* and *G. tsugae*, indicating that the ITS locus alone is limited in resolving species with confidence (Fig. S3). The limited power of ITS to differentiate species within the lucidum subclade is further demonstrated by the elevated machine-learning-based difficulty value for phylogenetic analyses (PyPythia V2. difficulty value = 0.57, Table S3, Fig. S4).

### The epitype highlights the paucity of knowledge about *Ganoderma lucidum* and clarifies its geographic distribution

The designation of a *G. lucidum* epitype provided an opportunity to revise our knowledge of this species; thus, we placed all publicly available *Ganoderma* genome assemblies, ITS sequences, and related metadata into this new context.

We inferred the most comprehensive phylogenomic tree of the genus *Ganoderma* to date, comprising 24 specimens, including seven *G. lucidum*-labelled genomes (Fig. 5/A, Fig. S5). The tree resolved three main clades corresponding to Subclades I, VII, and IX, with 100% ultrafast bootstrap support. Interestingly, none of the seven *G. lucidum*-labelled genomes clustered in subclade IX, where *G. lucidum* was expected; rather, all were located in subclade VII, which contains *G. lingzhi* and *G. resinaceum*. Next, we extracted seven barcoding loci from the genomes and placed them in a genus-level tree, which showed the same topology as the phylogenomic tree but included all subclades, providing further evidence that no *G. lucidum* genome has been sequenced to date.

**Figure 5.**
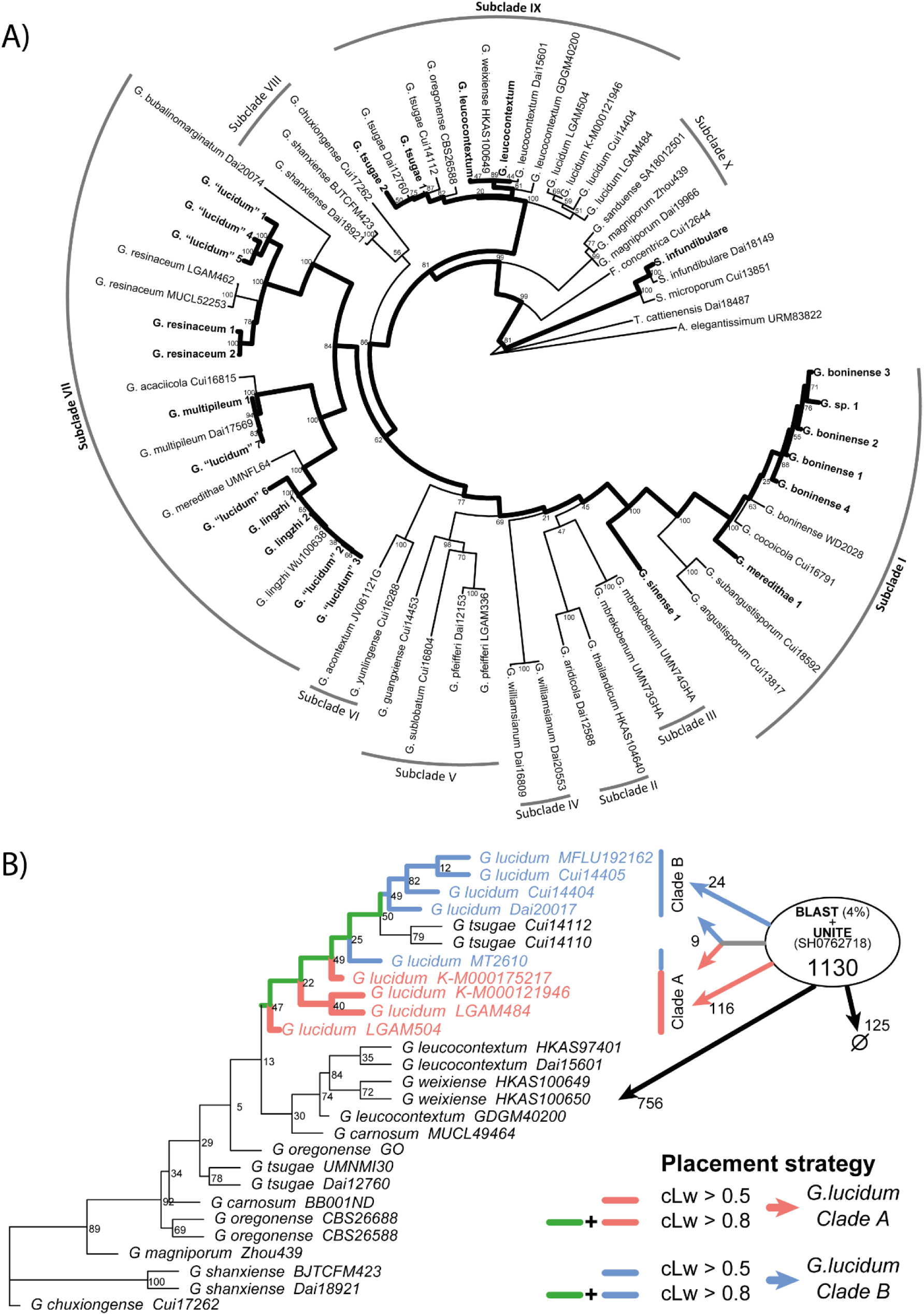
Genomic and barcoding knowledge gaps highlighted by the *G. lucidum* epitype. **A)** All available *Ganoderma* genomic data were placed in a genus-level tree showing that there is currently no sequenced genome of *G. lucidum* available. Bold branches highlight the topology of the phylogenomic tree. Note that seven genome assemblies are labelled as *G. lucidum*, but none of them clustered within the “lucidum” IX. clade. **B)** The results of the phylogenetic placement of potential *G. lucidum* sequences deposited in GenBank and UNITE databases. The colours used to label branches were matched to the placement strategy. cLw stands for cumulative Likelihood weight.

**Figure 6.**
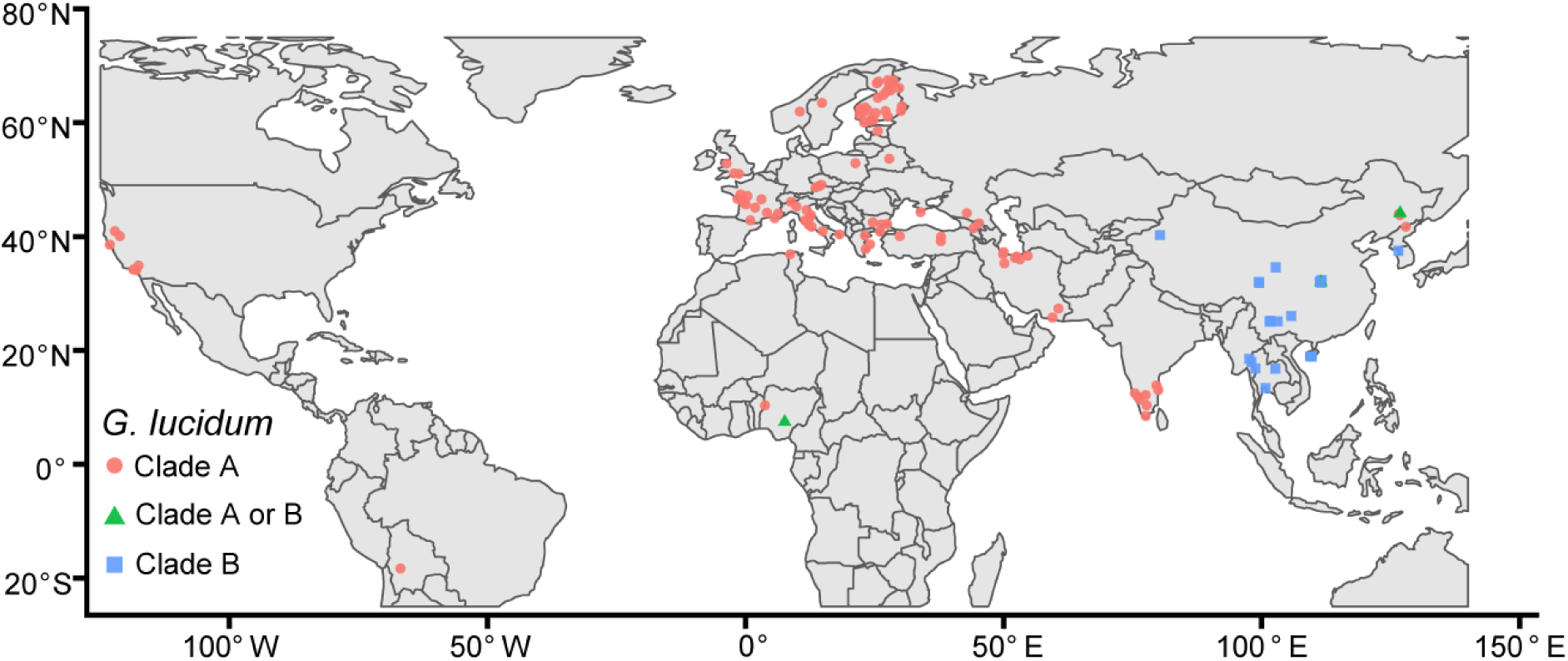
Geographic distribution of *G. lucidum* and its two clades based on the metadata of ITS sequences that are assigned to *G. lucidum* using phylogenetic placement with high support (cumulative Likelihood weight > 0.8).

To better understand the species’ distribution and available genetic information, we performed a phylogenetic placement of 1030 ITS sequences compiled from GenBank and UNITE databases. To do this, we found that the Gamma 4 model with LSU and SSU overhanging regions provided the most reliable inference compared to the species tree.

Because alignment thinning did not resolve the monophyly of *G. lucidum*, similar to the partitioned model above, we used the full tree to provide a more comprehensive backbone for phylogenetic placement (Figure 5/B). Out of the 1030 sequences, 149 were placed on branches linked to *G. lucidum* with high confidence (cLw > 0.8). All ITS sequences used to build the phylogenetic tree were assigned to the corresponding species, demonstrating the reliability of the phylogenetic placements. Interestingly, 749 sequences were labelled as *G. lucidum* in our compiled dataset, out of which only 128 were placed on the tree. This could be the result of the limitation of ITS barcoding demonstrated above, as well as short query sequences in the databases. The latter is supported by the fact that the sequence length distribution of unplaced *G. lucidum* sequences was 70 bp shorter on average compared to sequences that were identified as *G. lucidum*. We also separated the *G. lucidum* clade A and B sequences using phylogenetic placement, resulting in 116 and 24 sequences for Clade A and B, respectively, while 9 sequences couldn’t be distinguished between the two (Figure 5/B).

Finally, we used the 149 sequences confidently assigned to *G. lucidum* lineages to examine the geographic distribution and plant associations of this species (Table S4). Based on the 130 sequences with geographic information, *G. lucidum* shows a Eurasian distribution, with a few exceptions: six sequences were reported from North America (USA, California), one from South America (Bolivia), and three from Africa (Nigeria, Algeria). Within Eurasia, we found a clear geographic separation between clades A and B. The 20 Clade B sequences have an exclusive East Asian distribution, occurring in China and Thailand. Of the 110 Clade A sequences, four are also located in China, but one has no specific location, and the other three are labelled as *Polyporus tsugae* or as an uncultured fungus from Jilin and Changbai. The phylogenetic placement of these sequences was among the least confident (Table S4), suggesting that these instances might be misassigned to *G. lucidum*. Based on this, we have low support for an East Asian distribution of clade A. At the same time, we have strong evidence for a South Asian and Middle Eastern presence. Eight and nine specimens were reported from South India and Iran, respectively, some of which had a high cLw for Clade A branches only (cLW > 0.9).

Among the *G. lucidum* sequences assigned to Clade A, 34 harboured associated plant information at the genus or species level, from which 25 belonged to the order *Fagales*. With the exception of the epitype, none of these records originated from *Corylus*, but *Betulaceae* hosts accounted for eight records, including *Alnus*, *Betula* and *Carpinus*. Among *Fagaceae*, 16 records were associated with *Fagus*, *Castanea* and *Quercus*, the latter being the most frequent host genus (n = 14). Beyond *Fagales*, Clade A specimens were also reported from other hardwood genera, including *Malus*, *Platanus* and *Prunus*, as well as tropical and subtropical angiosperms such as *Azadirachta*, *Cassia*, *Tamarindus* and *Terminalia*. The East Asian Clade B showed a partly overlapping but distinct host spectrum. In addition to records from *Quercus*, specimens in this clade were also reported from fabaceous hosts (*Acacia*, *Pterocarpus*) and bamboo (*Dendrocalamus*).

## Discussion

Delineating and identifying economically and agronomically important species is crucial to fully harness their beneficial traits for human society (Pourkheirandish et al., 2020; Kweyamba et al., 2025; Davis et al., 2025). Medicinal mushrooms have been used in TCM for centuries, and recent preclinical and clinical studies have shown promising bioactive effects, including induction of neurogenesis and anticancer effects, among others (Powell, 2014). The most enigmatic and widely used medicinal mushroom is the “lingzhi” or “reishi” mushroom, which has been reported to have immunomodulatory, anticancer and anti-inflammatory effects (Wang et al., 2024b). It has a 6.2 billion USD market worldwide (www.grandviewresearch.com), yet the species identity has been debated for over two centuries, while the quantity and quality of medicinal compounds can vary widely across species and strains (Richter et al., 2015). The main sources of confusion stemmed from the lack of biological material served as a type for *Ganoderma lucidum*, the type species of the genus to which the “lingzhi” mushroom belongs. To resolve this, we designate here a sequenced interpretive specimen (i.e., epitype) to fix the application of the name *G. lucidum* 245 years after its first description (Curtis, 1781).

The selected specimen from the Kew Fungarium closely matches the original species concept in geographic origin, associated organism, and morphology, and provides a fixed reference point for future taxonomic and molecular studies. Our results further confirm that the true *G. lucidum* is distinct from a commonly conflated laccate species in the East Asian “lingzhi” clade VII, and other related European laccate *Ganoderma* species. Other micro-and macromorphological characteristics fit in the general concept of *G. lucidum* (Steyaert, 1967; Hennicke et al., 2016; Papp et al., 2017), further supporting the current morphological delineation of the species. Yet, due to the subtle micro-morphological differences and the high resemblance of closely related species to untrained eyes, molecular identification of *Ganoderma* species is essential.

We placed sequence data from the new epitype specimen in a genus-level time-calibrated tree, which was in line with the topology of recent phylogenies (Sun et al., 2022), and found that the “lucidum” clade was separated at least 12 million years ago from the “lingzhi” clade, suggesting vast evolutionary differences between the frequently conflated species. Moreover, the separation of the two lineages could have occurred even earlier, as other time calibrations found this group 5 to 15 Myr older (He et al., 2024; Szánthó et al., 2025; Varga et al., 2019).

By analysing the “lucidum” clade (Clade IX) at high resolution, we found that all six accepted species in the clade are resolved as monophyletic except for *G. weixiense* and *G. leucocontextum*, a pattern also found in previous studies (Fryssouli et al., 2020; Sun et al., 2022). We further detected two subclades within *G. lucidum* that have not been recognized previously: clade A harboured the epitype designated here, while clade B comprised five East Asian specimens.

Next, we assessed the usability of ITS barcoding for *Ganoderma lucidum* species delineation and identification, as this method is most commonly used in fungi and the ITS locus is the most abundant fungal molecular data employed in databases (Kauserud, 2023). We found that homology-based searches are unreliable, and searches of the GeneBank and UNITE databases contained 67% and 75% of sequences labelled as different species, respectively. This could stem from the low variability within the ITS loci of this group, which could be overcome to some extent through evolutionary modelling and by including SSU and LSU partial sequences overhanging the ITS sequence, but *G. lucidum* and *G. tsugae* still proved indistinguishable in many cases. ITS conservation within certain groups is known but rarely described, e.g. Garnica et al. (2016), which urges fungal evolutionary and ecological studies to use multi-locus approaches (Taylor et al., 2000) or at least to implement evolutionary modelling in sequence identification, such as phylogenetic placement methods (Kapli et al., 2017).

Consequently, we performed phylogenetic placement and assigned 149 ITS sequences to *G. lucidum* with high support, providing an opportunity to assess the species’ geographic distribution and associated plant preference. We found that *G. lucidum* has an Eurasian distribution with a few reported specimens from North and South America and Africa. It has already been suggested that *G. lucidum* was introduced to North America via mushroom farms (Loyd et al., 2018); and the South American or African sightings could suggest further intercontinental introduction. We further described the geographic isolation of Clades A and B, with Clade B occurring only in East Asia. Although the mPTP species delineation modelling did not distinguish these as separate species, geographic isolation could indicate cryptic speciation within *G. lucidum,* which would require more thorough morphological and genomic analyses. The associated organism data do not support a clear ecological distinction between Clades A and B, but suggest a broad spectrum of angiosperm wood for the whole lineage.

Finally, we found that the seven *G. lucidum* genomes published to date belong to *G. lingzhi*, *G. resinaceum,* and *G. multipileum*. Among these, the first *G*. “*lucidum*” genome (GCA_000271565 in Chen *et al*., 2012) was generated from the strain CGMCC5.0026 frequently used in commercial medicinal products. Yet, Chen et al. (2012) and other studies showing development-regulated chitin expression (Liu et al., 2025) or transcription factor-regulated triterpenoid synthesis (Li et al., 2026) involved species other than *G. lucidum*. On the other hand, a recent study treated *G. lucidum* and *G. lingzhi* as synonymous and analysed the genomes under a single name, yet, based on our analysis, only three of the seven belong to *G. lingzhi* (Wang et al., 2024a) (Figure 5).

The examples above underscore the importance of correctly identifying *G. lingzhi* and *G. lucidum* strains in genetic and pharmacological studies. Most pharmacological studies use triterpenoid extracts in which the compound composition is documented, e.g., (Das et al., 2020; Xia et al., 2020; Chen et al., 2023). Thus, based on these studies, we can link species’ health benefits with the quantity and the presence of certain compounds. Hennicke et al. (2016) measured the triterpenoid content of the *G. lucidum* M9720 strain, which was identified as clade A in our study, and found that its triterpenoid content was much lower than that of the *G. lingzhi* M9724 strain. Welti et al. (2015) did not detect *Ganoderma*-specific triterpenoids (Lucidenic acids and Ganoderic acids) in the *G. lucidum* strain CIRM-BRFM 885, which was also identified as clade A in our phylogenetic placement. The low triterpenoid content of *G. lucidum* suggests it may have different health benefits compared to *G. lingzhi*, but more thorough chemical profiling and preclinical tests of *G. lucidum* are needed to further clarify any possible health benefits. Thus, in future studies, molecular identification of strains used is essential, as triterpenoid content can vary substantially across species and strains.

Taken together, the separation of *G. lucidum*, *G. lingzhi*, and other laccate *Ganoderma* species is crucial for further advancing biodiversity, evolution, and drug discovery within this enigmatic medicinal mushroom group. Yet, conflating species and using both Latin and common names interchangeably remains common, despite the fact that *G. lucidum* is separated from the East Asian “lingzhi” species by more than 12 million years. Therefore, we propose “False lingzhi” as the English common name for *G*. *lucidum* and “Lingzhi” for *G*. *lingzhi* to support future studies and help policymakers and consumers communicate the distinction between the two commercially and culturally important species.

## Acknowledgement

TV and VP were supported by the Bentham Moxon Trust fund (BMT-V33-2024). The work of MGN was supported by the Fungarium Sequencing Project (FSP) via the Department for Environment, Food & Rural Affairs (DEFRA). The authors thank Lee Davies and the curators at Kew Fungarium for support in accessing the collection. We are grateful to Laszlo Csiba and Jehova Lourenco Junior for their help in the molecular lab and with scanning electron microscopy, respectively. We appreciate the help of the Fungarium Sequencing Project team at RBG, Kew, with DNA extraction. The authors acknowledge Research Computing at the James Hutton Institute for providing computational resources and technical support for the ‘UK’s Crop Diversity Bioinformatics HPC’ (BBSRC grants BB/S019669/1 and BB/X019683/1), use of which has contributed to the results reported within this paper. We appreciate the information about the *G. lucidum* sightings in the Peckham Rye area from Liam Nash & Clifford Davy of the Friends of Nunhead Cemetery. We thank Mohammad Sohrabi and Aida Sohrabi for their help in conducting a foray on 8 October 2025.

## Author contributions

TV, SRJP, and VP coordinated the work and prepared the initial draft. MGN performed the SEM analyses. VP and MGN conducted the morphological investigation. TV, MGN, SRJP and LH conducted the fieldwork. SRJP and LH carried out the molecular analyses. SRJP generated the preliminary results, and TV performed the final phylogenetic analyses. AA inferred the phylogenomic tree. AD and LH collected the georeferencing and host data, and AD visualised the spatial data. AMA and DH contributed to taxonomy and nomenclature. All authors participated in the study design and contributed to the writing and editing of the manuscript.

## Conflict of interest

The authors declare no conflict of interest.

## Data availability

Tree files, alignments and morphological measurements used in the study are attached as supplementary data. Newly generated sequences are deposited in the GeneBank database under the following accessions: PZ166757 and PZ179979.

among supplement files:

### Trees

Lucidum clade ML tree: LucidumPartitions_BS.treefile,

Genus-level ML tree: GenusPartitions_BS_rooted2.tree,

Dated genus tree: GenusTree_Dated.tre,

Genome tree: GenomeTree.treefile,

Genome-Barcode mixed tree: GenomeBarPartitions.treefile,

ITS tree for placements: LucidumITS_plus3_NoParG4.treefile

### Alignments

Genus-level ML tree: GenusSuperMatrix.fasta + GenusPartitions.txt.best_scheme.nex

Lucidum clade ML tree: LucidumSuperMatrix.fasta + LucidumPartitions.txt.best_scheme.nex

Genome tree: GenomeTree.faa + GenomeTree_best_scheme.nex

Genome-Barcode mixed tree: GenomBar_SuperMatrix.fasta +

GenomeBar_Partitions_best_scheme.nex

ITS tree for placements: LucidumTree_ITS_Muscle_trim.fasta

### Supplementary materials

**Supplementary Figure 1.** (SupplementaryFigure_1.pdf). The results of the mPTP species delineation analysis. Phylogenetic tree of the “lucidum subclade” with the *Gandoderma shanxiense* subclade as outgroup, in which branches are coloured on a gradient representing the posterior probability support of being a different speciation process (blacker) or the same speciation process (redder).

**Supplementary Figure 2.** (SupplementaryFigure_2.png). P-distance-based similarity comparison of species and specimens within the “lucidum subclade”.

**Supplementary Figure 3.** (SupplementaryFigure_3.pdf). ITS-based phylogenies were inferred using BaliPhy (Bayesian co-estimation of alignment and phylogeny) and IQ-tree (the top three phylogenies were reported).

**Supplementary Figure 4.** (SupplementaryFigure_4.pdf). Waterfall Plot depiction of Shapley values that shows which feature of the multisequence alignment contributes to the difficulty value estimated by PyPythia.

**Supplementary Figure 5.** (SupplementaryFigure_5.pdf). Maximum likelihood phylogeny based on all available *Ganoderma* genomes.

**Supplementary Table 1.** (SupplementaryTable_1.xlsx). List of specimens and related genomic and genetic data used to infer genus-level tree, the tree of subclade 9 (“lucidum subclade”), the phylogenomic tree and the genus-level tree where sequences were also retrieved from genome assemblies.

**Supplementary Table 2.** (SupplementaryTable_2.xlsx). Raw spore measurements of the epirype specimen.

**Supplementary Table 3.** (SupplementaryTable_3.xlsx). Comparison of multisequence alignment-based difficulty value and other parameters of different loci within the “lucidum subclade”.

**Supplementary Table 4.** (SupplementaryTable_4.xlsx). The results of the phylogenetic placement.

## Notes

### Competing Interest Statement

The authors have declared no competing interest.

## References

Abarenkov, K., Nilsson, R. H., Larsson, K.-H., Taylor, A. F., May, T. W., Frøslev, T. G., Pawlowska, J., Lindahl, B., Põldmaa, K., & Truong, C. (2024). The UNITE database for molecular identification and taxonomic communication of fungi and other eukaryotes: sequences, taxa and classifications reconsidered. Nucleic Acids Research, 52(D1), D791–D797.

Altschul, S. F., Gish, W., Miller, W., Myers, E. W., & Lipman, D. J. (1990). Basic local alignment search tool. Journal of molecular biology, 215(3), 403–410.

Barbera, P., Kozlov, A. M., Czech, L., Morel, B., Darriba, D., Flouri, T., & Stamatakis, A. (2019). EPA-ng: massively parallel evolutionary placement of genetic sequences. Systematic biology, 68(2), 365–369.

Barua, R. C., Coniglio, R. O., Molina, M. A., Díaz, G. V., & Fonseca, M. I. (2024). Fungi as biotechnological allies: Exploring contributions of edible and medicinal mushrooms. Journal of Food Science, 89(11), 6888–6915.

Bengtsson-Palme, J., Ryberg, M., Hartmann, M., Branco, S., Wang, Z., Godhe, A., De Wit, P., Sánchez-García, M., Ebersberger, I., & De Sousa, F. (2013). Improved software detection and extraction of ITS1 and ITS2 from ribosomal ITS sequences of fungi and other eukaryotes for analysis of environmental sequencing data. Methods in Ecology and Evolution, 914–919.

Bennett, D. J., Hettling, H., Silvestro, D., Vos, R., & Antonelli, A. (2018). restez: Create and Query a Local Copy of GenBank in R. Journal of Open Source Software, 3(31), 1102.

Bhunjun, C., Chen, Y., Phukhamsakda, C., Boekhout, T., Groenewald, J., McKenzie, E., Francisco, E., Frisvad, J., Groenewald, M., Hurdeal, V., Luangsa-Ard, J., Perrone, G., Visagie, C. M., Bai, F. Y., Błaszkowski, J., Braun, U., de Souza, F. A., de Queiroz, M. B., Dutta, A. K. … Crous, P. W. (2024). What are the 100 most cited fungal genera? Studies in Mycology, 108(1), 1–412.

Bishop, K. S., Kao, C. H., Xu, Y., Glucina, M. P., Paterson, R. R. M., & Ferguson, L. R. (2015). From 2000 years of *Ganoderma lucidum* to recent developments in nutraceuticals. Phytochemistry, 114, 56–65.

Borowiec, M. L. (2016). AMAS: a fast tool for alignment manipulation and computing of summary statistics. PeerJ, 4, e1660.

Bradshaw, M. J., Aime, M. C., Rokas, A., Maust, A., Moparthi, S., Jellings, K., Pane, A. M., Hendricks, D., Pandey, B., Li, Y., & Pfister, D. H. (2023). Extensive intragenomic variation in the internal transcribed spacer region of fungi. iScience, 26(8), 107317.

Cao, Y., Wu, S.-H., & Dai, Y.-C. (2012). Species clarification of the prize medicinal *Ganoderma* mushroom “Lingzhi”. Fungal Diversity, 56(1), 49–62.

Capella-Gutiérrez, S., Silla-Martínez, J. M., & Gabaldón, T. (2009). trimAl: a tool for automated alignment trimming in large-scale phylogenetic analyses. Bioinformatics, 25(15), 1972–1973.

Charif, D., & Lobry, J. R. (2007). SeqinR 1.0-2: A Contributed Package to the R Project for Statistical Computing Devoted to Biological Sequences Retrieval and Analysis, in Bastolla, U., Porto, M., Roman, H.E., & Vendruscolo, M. (eds.) Structural Approaches to Sequence Evolution: Molecules, Networks, Populations. Berlin, Heidelberg: Springer Berlin Heidelberg, 207–232.

Chen, S.-N., Nan, F.-H., Liu, M.-W., Yang, M.-F., Chang, Y.-C., & Chen, S. (2023). Evaluation of immune modulation by β-1,3; 1,6 D-glucan derived from *Ganoderma lucidum* in healthy adult volunteers, a randomized controlled trial. Foods, 12(3), 659.

Chen, S., Xu, J., Liu, C., Zhu, Y., Nelson, D. R., Zhou, S., Li, C., Wang, L., Guo, X., & Sun, Y. (2012). Genome sequence of the model medicinal mushroom *Ganoderma lucidum*. Nature communications, 3(1), 913.

Cortina-Escribano, M., Veteli, P., Wingfield, M. J., Wingfield, B. D., Coetzee, M. P. A., Vanhanen, H., & Linnakoski, R. (2024). Phylogenetic analysis and morphological characteristics of laccate *Ganoderma* specimens in Finland. Mycologia, 116(6), 1046–1062.

Curtis, W. (1781) Flora Londinensis. London: Printed for and sold by the author and B. White, 71.

Dai, Y.-C., Zhou, L.-W., Hattori, T., Cao, Y., Stalpers, J. A., Ryvarden, L., Buchanan, P., Oberwinkler, F., Hallenberg, N., & Liu, P.-G. (2017). *Ganoderma lingzhi* (Polyporales, Basidiomycota): the scientific binomial for the widely cultivated medicinal fungus Lingzhi’, Mycological Progress, 16(11), 1051–1055.

Das, A., Alshareef, M., Henderson Jr, F., Martinez Santos, J., Vandergrift III, W., Lindhorst, S. M., Varma, A. K., Infinger, L., Patel, S. J., & Cachia, D. (2020). Ganoderic acid A/DM-induced NDRG2 over-expression suppresses high-grade meningioma growth. Clinical and Translational Oncology, 22(7), 1138–1145.

Davis, A. P., Shepherd-Clowes, A., Cheek, M., Moat, J., Wei Luo, D., Kiwuka, C., Kalema, J., Tchiengué, B., & Viruel, J. (2025). Genomic data define species delimitation in *Liberica* coffee with implications for crop development and conservation. Nature Plants, 11(9), 1729–1738.

Du, Y., Tian, L., Wang, Y., Li, Z., & Xu, Z. (2024). Chemodiversity, pharmacological activity, and biosynthesis of specialized metabolites from medicinal model fungi *Ganoderma lucidum*. Chinese Medicine, 19(1), 51.

Du, Z., Li, Y., Wang, X.-C., Wang, K., & Yao, Y.-J. (2023). Re-examination of the holotype of *Ganoderma sichuanense* (Ganodermataceae, Polyporales) and a clarification of the identity of Chinese cultivated lingzhi. Journal of Fungi, 9(3), 323.

Edgar, R. C. (2022). Muscle5: High-accuracy alignment ensembles enable unbiased assessments of sequence homology and phylogeny. Nature communications, 13(1), 6968.

Fleischmann, A., Krings, M., Mayr, H., & Agerer, R. (2007). Structurally preserved polypores from the Neogene of North Africa: *Ganodermites libycus gen. et sp. nov.* (Polyporales, Ganodermataceae). Review of Palaeobotany and Palynology, 145(1–2), 159–172.

Fryssouli, V., Zervakis, G. I., Polemis, E., & Typas, M. A. (2020). A global meta-analysis of ITS rDNA sequences from material belonging to the genus *Ganoderma* (Basidiomycota, Polyporales) including new data from selected taxa. MycoKeys, 75, 71.

Garnica, S., Schön, M. E., Abarenkov, K., Riess, K., Liimatainen, K., Niskanen, T., Dima, B., Soop, K., Frøslev, T. G., & Jeppesen, T. S. (2016). Determining threshold values for barcoding fungi: lessons from *Cortinarius* (Basidiomycota), a highly diverse and widespread ectomycorrhizal genus. FEMS Microbiology Ecology, 92(4), fiw045.

Haag, J., Höhler, D., Bettisworth, B., & Stamatakis, A. (2022). From easy to hopeless-predicting the difficulty of phylogenetic analyses. Molecular Biology and Evolution, 39(12), msac254.

He, M.-Q., Cao, B., Liu, F., Boekhout, T., Denchev, T. T., Schoutteten, N., Denchev, C. M., Kemler, M., Gorjón, S. P., Begerow, D., Valenzuela, R., Davoodian, N., Niskanen, T., Vizzini, A., Redhead, S. A., Ramírez-Cruz, V., Papp, V., Dudka, V. A., Dutta, A. K., … Zhao, R.-L. (2024). Phylogenomics, divergence times and notes of orders in Basidiomycota. Fungal diversity, 126(1), 27–406.

Heibl, C. (2008). PHYLOCH: R language tree plotting tools and interfaces to diverse phylogenetic software packages. Retrieved from http://www.christophheibl.de/Rpackages.html

Hennicke, F., Cheikh-Ali, Z., Liebisch, T., Maciá-Vicente, J. G., Bode, H. B., & Piepenbring, M. (2016). Distinguishing commercially grown *Ganoderma lucidum* from *Ganoderma lingzhi* from Europe and East Asia on the basis of morphology, molecular phylogeny, and triterpenic acid profiles. Phytochemistry, 127, 29–37.

Hibbett, D., Abarenkov, K., Kõljalg, U., Öpik, M., Chai, B., Cole, J., Wang, Q., Crous, P., Robert, V., Helgason, T., Herr, J. R., Kirk, P., Lueschow, S., O’Donnell, K., Nilsson, R. H., Oono, R., Schoch, C., Smyth, C., Walker, D.M., … Geiser, D. M. (2016). Sequence-based classification and identification of Fungi. Mycologia, 108(6), 1049–1068.

Hill, R. (2021) GenePull. Available at: https://github.com/Rowena-h/MiscGenomicsTools/tree/main/GenePull.

Hill, R. (2024). BUSCOfilter. Available at: https://github.com/Rowena-h/BUSCOfilter.

Hoang, D. T., Chernomor, O., von Haeseler, A., Minh, B. Q. & Vinh, L. S. (2018). UFBoot2: improving the ultrafast bootstrap approximation. Molecular Biology and Evolution, 35, 518–522.

Hodgson, E. & Varga, T. (2023). PhyloGenie. Available at: https://github.com/vtorda/PhyloGenie.

Jafari, A., Mardani, H., Mirzaei Fashtali, Z., & Arghavan, B. (2025). The Nutritional Significance of *Ganoderma lucidum* on Human Health: A GRADE-Assessed Systematic Review and Meta-Analysis of Clinical Trials. Food science & nutrition, 13(6), e70423.

Kalyaanamoorthy, S., Minh, B., Wong, T., von Haeseler, A., & Jermiin, L. S. (2017). ModelFinder: fast model selection for accurate phylogenetic estimates. Nature Methods, 14, 587–589.

Kapli, P., Lutteropp, S., Zhang, J., Kobert, K., Pavlidis, P., Stamatakis, A., & Flouri, T. (2017). Multi-rate Poisson tree processes for single-locus species delimitation under maximum likelihood and Markov chain Monte Carlo. Bioinformatics, 33(11), 1630–1638.

Karunarathna, S. C., Prasannath, K., Lu, W., & Hapuarachchi, K. K. (2025). *Ganoderma*: bridging traditional wisdom with modern innovation in medicinal mushroom and dietary supplement industry. New Zealand Journal of Botany, 63(5), 2529–2588.

Katoh, K., & Standley, D. M. (2013). MAFFT Multiple Sequence Alignment Software Version 7: Improvements in Performance and Usability. Molecular Biology and Evolution, 30(4), 772–780.

Kauserud, H. (2023). ITS alchemy: On the use of ITS as a DNA marker in fungal ecology. Fungal Ecology, 65, 101274.

Klupp, N. L., Kiat, H., Bensoussan, A., Steiner, G. Z., & Chang, D. H. (2016). A double-blind, randomised, placebo-controlled trial of *Ganoderma lucidum* for the treatment of cardiovascular risk factors of metabolic syndrome. Scientific reports, 6(1), 29540.

Kõljalg, U., Abarenkov, K., Nilsson, R. H., Larsson, K.-H., May, T. W., Taylor, A. F. S., Frøslev, T. G., & Põldmaa, K. (2024) ’TH100067: Ganoderma’. Available at: 10.15156/BIO/TH100067 (Accessed.

Kweyamba, P. A., Hofer, L. M., Kibondo, U. A., Mwanga, R. Y., Sayi, R. M., Matwewe, F., Lwetoijera, D. W., Tambwe, M. M., & Moore, S. J. (2025). Contrasting vector competence of three main East African Anopheles malaria vector mosquitoes for *Plasmodium falciparum*. Scientific Reports 15(1), 2286.

Larsson, A. (2014). AliView: a fast and lightweight alignment viewer and editor for large datasets. Bioinformatics, 30(22), 3276–3278.

Latorre, S. M., Lang, P. L., Burbano, H. A., & Gutaker, R. M. (2020). Isolation, library preparation, and bioinformatic analysis of historical and ancient plant DNA. Current Protocols in Plant Biology, 5(4), e20121.

Li, Y., Xu, L., Cui, J., Wang, S., Yuan, Y., Xiao, C., Luo, X., Baleev, D., Zhan, Y., & Yin, J. (2026). Analysis of BpbHLH Gene Family Responsive to MeJA Signalling in *Betula platyphylla* Suk. and Functional Mechanisms of BpbHLH42/44 in Genetic Improvement and Triterpenoid Biosynthesis. Plant Biotechnology Journal, 10.1111/pbi.70626

Liang, C., Tian, D., Liu, Y., Li, H., Zhu, J., Li, M., Xin, M., & Xia, J. (2019). Review of the molecular mechanisms of *Ganoderma lucidum* triterpenoids: Ganoderic acids A, C2, D, F, DM, X and Y. European Journal of Medicinal Chemistry, 174, 130–141.

Liu, L., Yang, Y., Li, J., Gao, Y., & Yan, M. (2025). Analysis of the chitin synthase gene family in *Ganoderma lucidum*: its structure, phylogeny, and expression patterns. PeerJ, 13, e20302.

Loyd, A. L., Richter, B. S., Jusino, M. A., Truong, C., Smith, M. E., Blanchette, R. A., & Smith, J. A. (2018). Identifying the “mushroom of immortality”: assessing the *Ganoderma* species composition in commercial Reishi products. Frontiers in Microbiology, 9, 1557.

Löytynoja, A. (2013). Phylogeny-aware alignment with PRANK. Multiple sequence alignment methods: Springer, 155–170.

Matsen, F. A., Kodner, R. B., & Armbrust, E. V. (2010). pplacer: linear time maximum-likelihood and Bayesian phylogenetic placement of sequences onto a fixed reference tree. BMC bioinformatics, 11(1), 538.

Moncalvo, J.-M., & Ryvarden, L. (1997). A nomenclatural study of the Ganodermataceae Donk. Synopsis Fungorum, 11, 1–114.

Moncalvo, J.-M., Wang, H.-F., & Hseu, R.-S. (1995a). Gene phylogeny of the *Ganoderma lucidum* complex based on ribosomal DNA sequences. Comparison with traditional taxonomic characters. Mycological research, 99(12), 1489–1499.

Moncalvo, J.-M., Wang, H.-H., & Hseu, R.-S. (1995b). Phylogenetic relationships in *Ganoderma* inferred from the internal transcribed spacers and 25S ribosomal DNA sequences. Mycologia, 87(2), 223–238.

Pagès, H., Aboyoun, P., Gentleman, R., & DebRoy, S. (2021). Biostrings: efficient manipulation of biological strings R package version 2.62.0.: Bioconductor.

Papp, V. (2019). Global Diversity of the Genus *Ganoderma* Taxonomic Uncertainties and Challenges. Advances in Macrofungi: CRC Press, pp. 10–33.

Papp, V. (2024). The Lingzhi naming dilemma: Overlooked and long-forgotten names threaten nomenclatural stability. Fungal Biology Reviews, 47, 100338.

Papp, V., Dima, B., & Wasser, S. P. (2017). What is *Ganoderma lucidum* in the molecular era? International Journal of Medicinal Mushrooms, 19(7), 575–593.

Paradis, E., & Schliep, K. (2019). ape 5.0: an environment for modern phylogenetics and evolutionary analyses in R. Bioinformatics, 35(3), 526–528.

Paterson, R. R. M. (2006). *Ganoderma*–a therapeutic fungal biofactory. Phytochemistry, 67(18), 1985–2001.

Pourkheirandish, M., Golicz, A. A., Bhalla, P. L., & Singh, M. B. (2020). Global role of crop genomics in the face of climate change. Frontiers in plant science, 11, 922.

Powell, M. (2014). Medicinal mushrooms: a clinical guide. 2nd edition. edn. Eastbourne, East Sussex, UK: Mycology Press, an imprint of Bamboo Publishing Ltd.

Redelings, B. D. (2021). BAli-Phy version 3: model-based co-estimation of alignment and phylogeny. Bioinformatics, 37(18), 3032–3034.

Redelings, B. D., & Suchard, M. A. (2007). Incorporating indel information into phylogeny estimation for rapidly emerging pathogens. BMC evolutionary biology, 7(1), 40.

Richter, C., Wittstein, K., Kirk, P. M., & Stadler, M. (2015). An assessment of the taxonomy and chemotaxonomy of *Ganoderma*. Fungal Diversity, 71(1), 1–15.

Schliep, K. P. (2011). phangorn: phylogenetic analysis in R. Bioinformatics, 27(4), 592–593.

Schoch, C. L., Seifert, K. A., Huhndorf, S., Robert, V., Spouge, J. L., Levesque, C. A., Chen, W., & Fungal Barcoding Consortium (2012). Nuclear ribosomal internal transcribed spacer (ITS) region as a universal DNA barcode marker for Fungi. Proceedings of the National Academy of Sciences, 109(16), 6241–6246.

Smith, S. A., & O’Meara, B. C. (2012). treePL: divergence time estimation using penalized likelihood for large phylogenies. Bioinformatics, 28(20), 2689–2690.

South, A., Michael, S., & Massicotte, P. (2026). rnaturalearthdata: world vector map data from Natural Earth used in ’rnaturalearth’. CRAN: Contributed Packages.

Steenwyk, J. L., Buida, T. J. III, Li, Y., Shen, X. X., & Rokas, A. (2020). ClipKIT: A multiple sequence alignment trimming software for accurate phylogenomic inference. PLoS Biology 18(12), e3001007.

Stevenson, A., Dandy, J. E., & Stearn, W. T. (1961). A bibliographic study of William Curtis’ Flora Londinensis 1777–98[1775–]. Volume II Catalogue of botanical books in the collection of Rachel McMasters Miller Hunt. Pittsburgh, PA, USA: Hunt Foundation.

Steyaert, R. L. (1961). Note on the nomenclature of fungi and, incidentally, of *Ganoderma lucidum*. Taxon, 10(8), 251.

Steyaert, R. L. (1967). Considérations générales sur le genre *Ganoderma* et plus spécialement sur les espèces européennes. Bulletin de la Société Royale de Botanique de Belgique/Bulletin van de Koninklijke Belgische Botanische Vereniging, 189–211.

Sun, Y.-F., Xing, J., He, X., Wu, D., Song, C., Liu, S., Vlasák, J., Gates, G., Gibertoni, T., & Cui, B. (2022). Species diversity, systematic revision and molecular phylogeny of Ganodermataceae (Polyporales, Basidiomycota) with an emphasis on Chinese collections. Studies in Mycology, 101(1), 287–415.

Szánthó, L. L., Merényi, Z., Donoghue, P., Gabaldón, T., Nagy, L. G., Szöllősi, G. J., & Ocaña-Pallarès, E. (2025). A timetree of Fungi dated with fossils and horizontal gene transfers. Nature Ecology & Evolution, 9(11), 1989–2001.

Szedlay, G. (2002). Is the widely used medicinal fungus the *Ganoderma lucidum* (Fr.) Karst. sensu stricto? Acta Microbiologica et Immunologica Hungarica, 49(2–3), 235–243.

Taylor, J. W. (2011). One Fungus = One Name: DNA and fungal nomenclature twenty years after PCR. IMA Fungus, 2(2), 113–120.

Taylor, J. W., Jacobson, D. J., Kroken, S., Kasuga, T., Geiser, D. M., Hibbett, D. S., & Fisher, M. C. (2000). Phylogenetic species recognition and species concepts in fungi. Fungal genetics and biology, 31(1), 21–32.

Tegenfeldt, F., Kuznetsov, D., Manni, M., Berkeley, M., Zdobnov, E. M., & Kriventseva, E. V. (2025). OrthoDB and BUSCO update: annotation of orthologs with wider sampling of genomes. Nucleic acids research, 53(D1), D516–D522.

Thomson, S. A., Pyle, R. L., Ahyong, S. T., Alonso-Zarazaga, M., Ammirati, J., Araya, J. F., Ascher, J. S., Audisio, T. L., Azevedo-Santos, V. M., Bailly, N., Baker, W. J., Balke, M., Barclay, M. V. L., Barrett, R. L., Benine, R. C., Bickerstaff, J. R. M., Bouchard, P., Bour, R., Bourgoin, T., … Zhou, H.-Z. (2018). Taxonomy based on science is necessary for global conservation. PLoS biology, 16(3), e2005075.

Turland, N. J., Wiersema, J. H., Barrie, F. R., Gandhi, K. N., Gravendyck, J., Greuter, W., Hawksworth, D. L., Herendeen, P. S., Klopper, R. R., Knapp, S., Kusber, W.-H., Li, D.-Z., May, T. W., Monro, A. M., Prado, J., Price, M. J., Smith, G. F., & Zamora Señoret, J.C. (2025). International Code of Nomenclature for algae, fungi, and plants (Madrid Code). Regnum Vegetabile 162. Chicago: University of Chicago Press. 10.7208/chicago/9780226839479.001.0001

Varga, T., Krizsán, K., Földi, C., Dima, B., Sánchez-García, M., Sánchez-Ramírez, S., Szöllősi, G. J., Szarkándi, J. G., Papp, V., Albert, L., Andreopoulos, W., Angelini, C., Antonín, V., Barry, K. W., Bougher, N. L., Buchanan, P., Buyck, B., Bense, V., Catcheside, P., … Nagy, L. G. (2019). Megaphylogeny resolves global patterns of mushroom evolution. Nature ecology & evolution, 3(4), 668–678.

Wang, D.-M., Wu, S.-H., Su, C.-H., Peng, J.-T., Shih, Y.-H., & Chen, L.-C. (2009). *Ganoderma multipileum*, the correct name for ‘*G. lucidum*’ in tropical Asia. Botanical studies, 50(4), 451–458.

Wang, L., Shi, P., Ping, Z., Huang, Q., Jiang, L., Ma, N., Wang, Q., Xu, J., Zou, Y., & Huang, Z. (2024a). The golden genome annotation of *Ganoderma lingzhi* reveals a more complex scenario of eukaryotic gene structure and transcription activity. BMC biology, 22(1), 271.

Wang, S., Wang, L., Shangguan, J., Jiang, A., & Ren, A. (2024b). Research progress on the biological activity of ganoderic acids in *Ganoderma lucidum* over the last five years. Life, 14(10), 1339.

Wang, X.-C., Xi, R.-J., Li, Y., Wang, D.-M., & Yao, Y.-J. (2012). The species identity of the widely cultivated *Ganoderma*, ‘*G. lucidum*’ (Ling-zhi), in China. Plos one, 7(7), e40857.

Welti, S., Moreau, P. A., Decock, C., Danel, C., Duhal, N., Favel, A., & Courtecuisse, R. (2015). Oxygenated lanostane-type triterpenes profiling in laccate *Ganoderma* chemotaxonomy. Mycological Progress, 14(7), 45.

White, T. J., Bruns, T., Lee, S., & Taylor, J. (1990). Amplification and direct sequencing of fungal ribosomal RNA genes for phylogenetics. PCR protocols: a guide to methods and applications, 18(1), 315–322.

Wickham, H. (2009). ggplot2 Elegant graphics for data analysis introduction. Use R. Springer-Verlag, 1007, 978–0.

Winter, D. J. (2017). Rentrez: An R Package for the NCBI eUtils API. PeerJ Preprints, 5, e3179v2.

Wong, T. K., Ly-Trong, N., Ren, H., Baños, H., Roger, A. J., Susko, E., Bielow, C., De Maio, N., Goldman, N., Hahn, M. W., dos Reis, M., Vinh, L. S., Huttley, G., Lanfear, R., Minh, B. Q. (2026). IQ-TREE 3: phylogenomic inference software using complex evolutionary models. Molecular Biology and Evolution, msag117, 10.1093/molbev/msag117

Wu, S., Zhang, S., Peng, B., Tan, D., Wu, M., Wei, J., Wang, Y., & Luo, H. (2024). *Ganoderma lucidum*: a comprehensive review of phytochemistry, efficacy, safety and clinical study. Food Science and Human Wellness, 13(2), 568–596.

Xia, J., Dai, L., Wang, L., & Zhu, J. (2020). Ganoderic acid DM induces autophagic apoptosis in non-small cell lung cancer cells by inhibiting the PI3K/Akt/mTOR activity’, Chemico-Biological Interactions, 316, 108932.

Yao, Y.-J., Li, Y., Du, Z., Wang, K., Wang, X.-C., Kirk, P. M., & Spooner, B. M. (2020). On the typification of *Ganoderma sichuanense* (Agaricomycetes)−the widely cultivated lingzhi medicinal mushroom. International journal of medicinal mushrooms, 22(1), 45–54.

Yao, Y.-J., Wang, X.-C., & Wang, B. (2013). Epitypification of *Ganoderma sichuanense* JD Zhao & XQ Zhang (Ganodermataceae). Taxon, 62(5), 1025–1031.

Zhou, L.-W., Cao, Y., Wu, S.-H., Vlasák, J., Li, D.-W., Li, M.-J., & Dai, Y.-C. (2015). Global diversity of the *Ganoderma lucidum* complex (Ganodermataceae, Polyporales) inferred from morphology and multilocus phylogeny. Phytochemistry, 114, 7–15.

