## Supplementary material for "A historical specimen of False Lingzhi (*Ganoderma lucidum*) resolves a 245-year-old confusion within an important medicinal mushroom group": SupplementaryFigure_3.pdf

IQTree: ITS+SSU,LSU overhanging + RPB2, partitioned

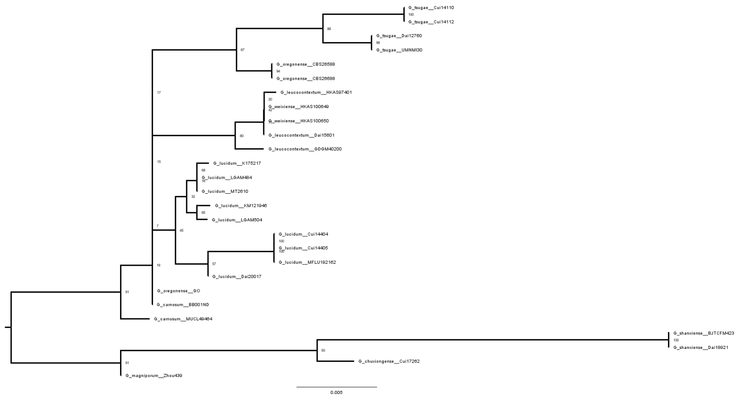

BaliPhy: ITS, Partitioned

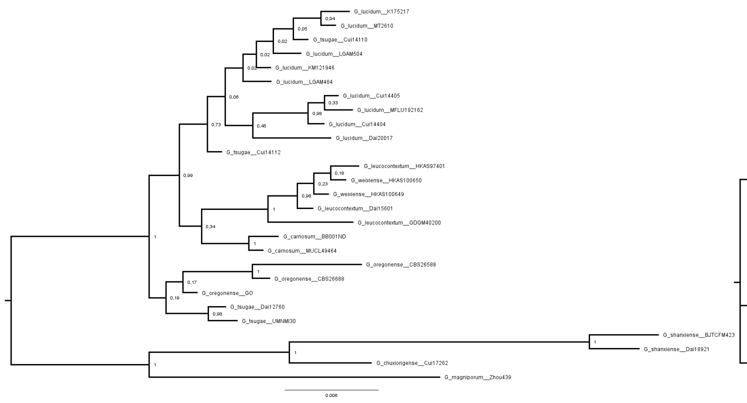

BaliPhy: ITS thinned, Partitioned

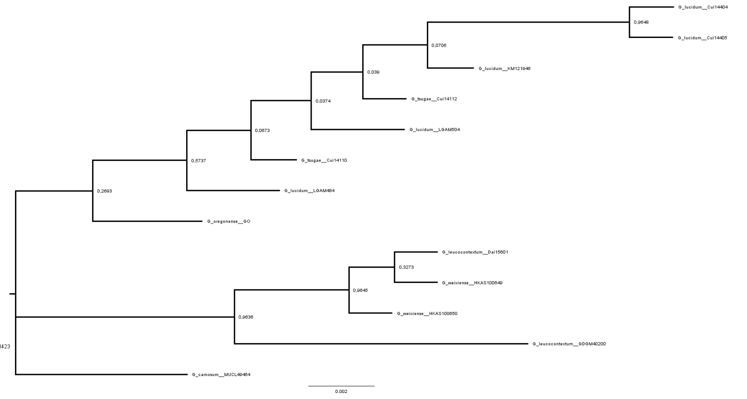

IQTree: ITS+SSU,LSU overhanging, Partitioned

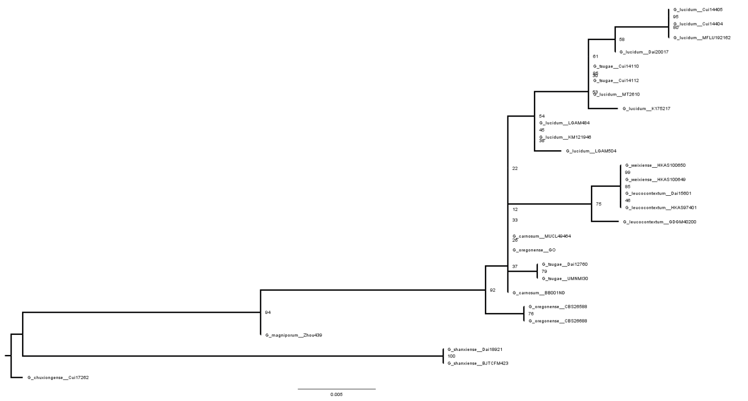

BaliPhy: ITS+SSU,LSU overhanging, Not-Partitioned

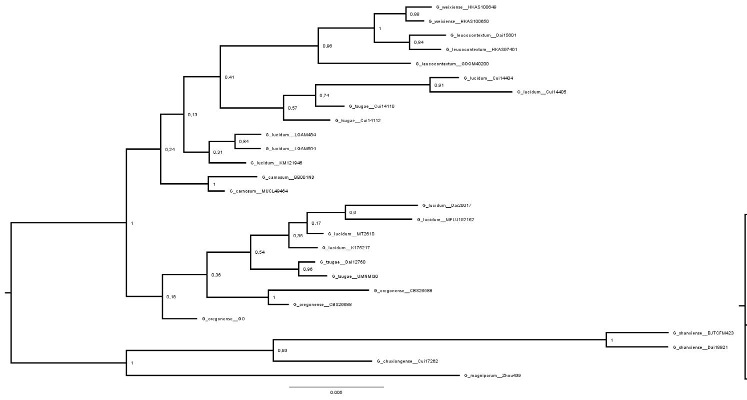

BaliPhy: ITS thinned, Not-Partitioned

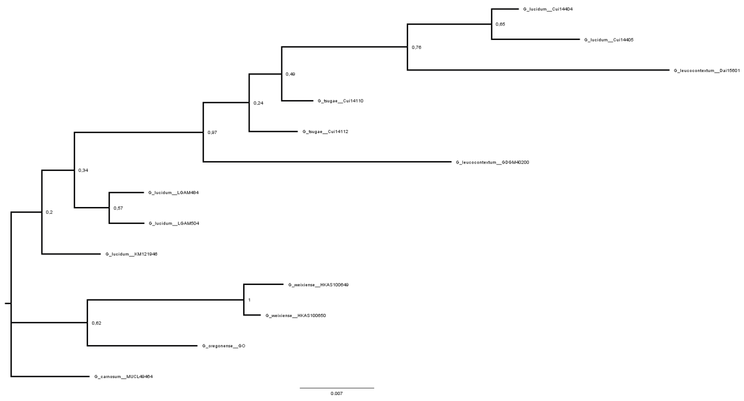

IQTree: ITS+SSU,LSU overhanging, thinned Partitioned

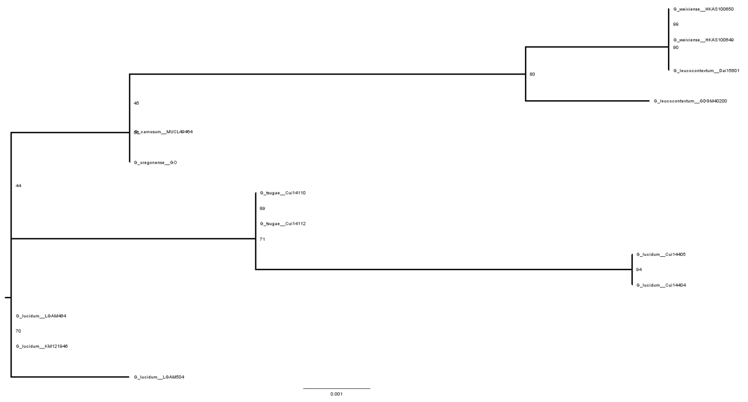

BaliPhy: ITS+SSU,LSU overhanging, Partitioned

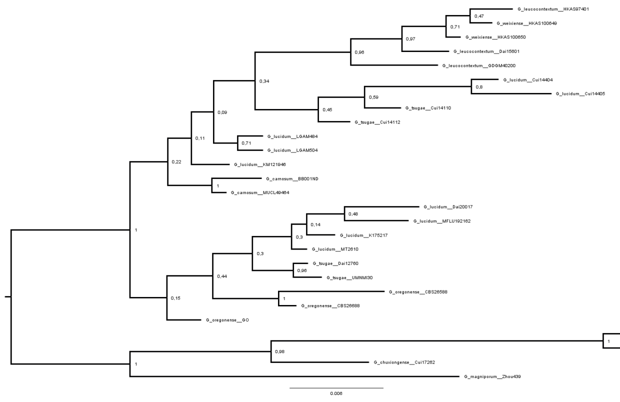

BaliPhy: ITS+SSU,LSU overhanging, thinned, Partitioned

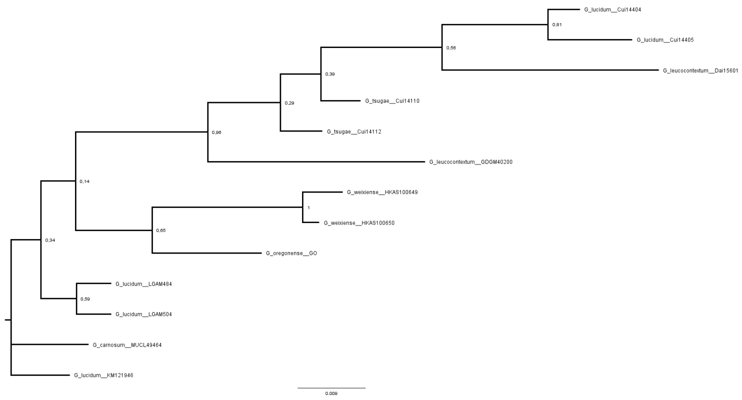
