## Supplementary figures and images for "A historical specimen of False Lingzhi (*Ganoderma lucidum*) resolves a 245-year-old confusion within an important medicinal mushroom group"

### SupplementaryFigure_1.pdf

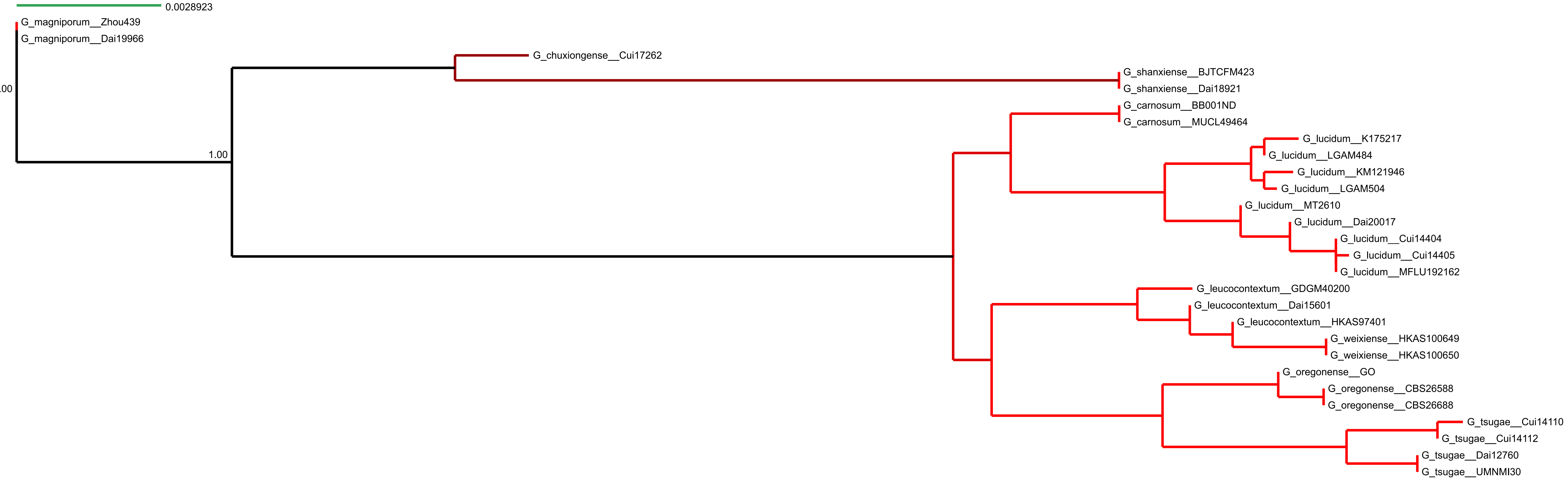

### SupplementaryFigure_2.png

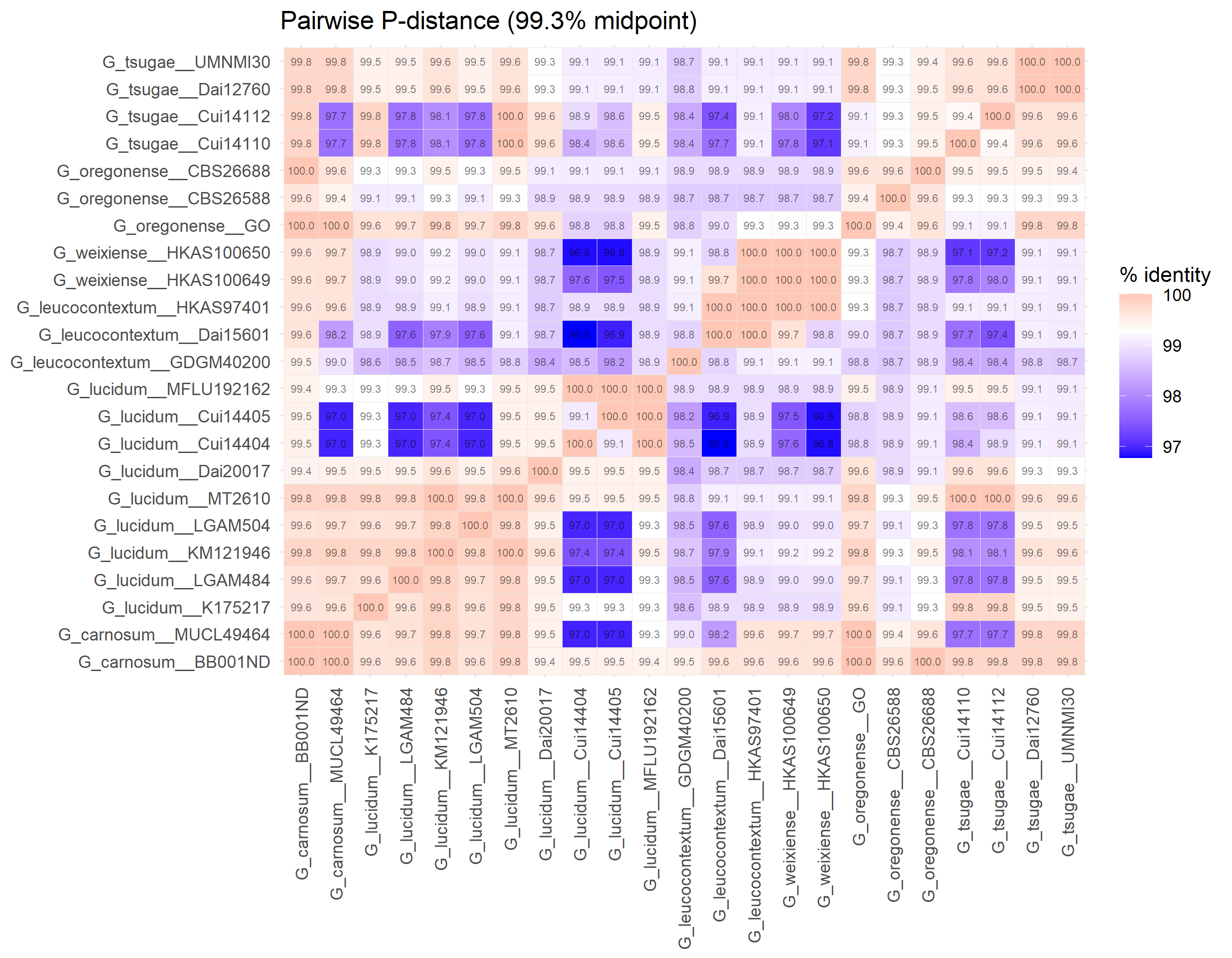

### SupplementaryFigure_4.pdf

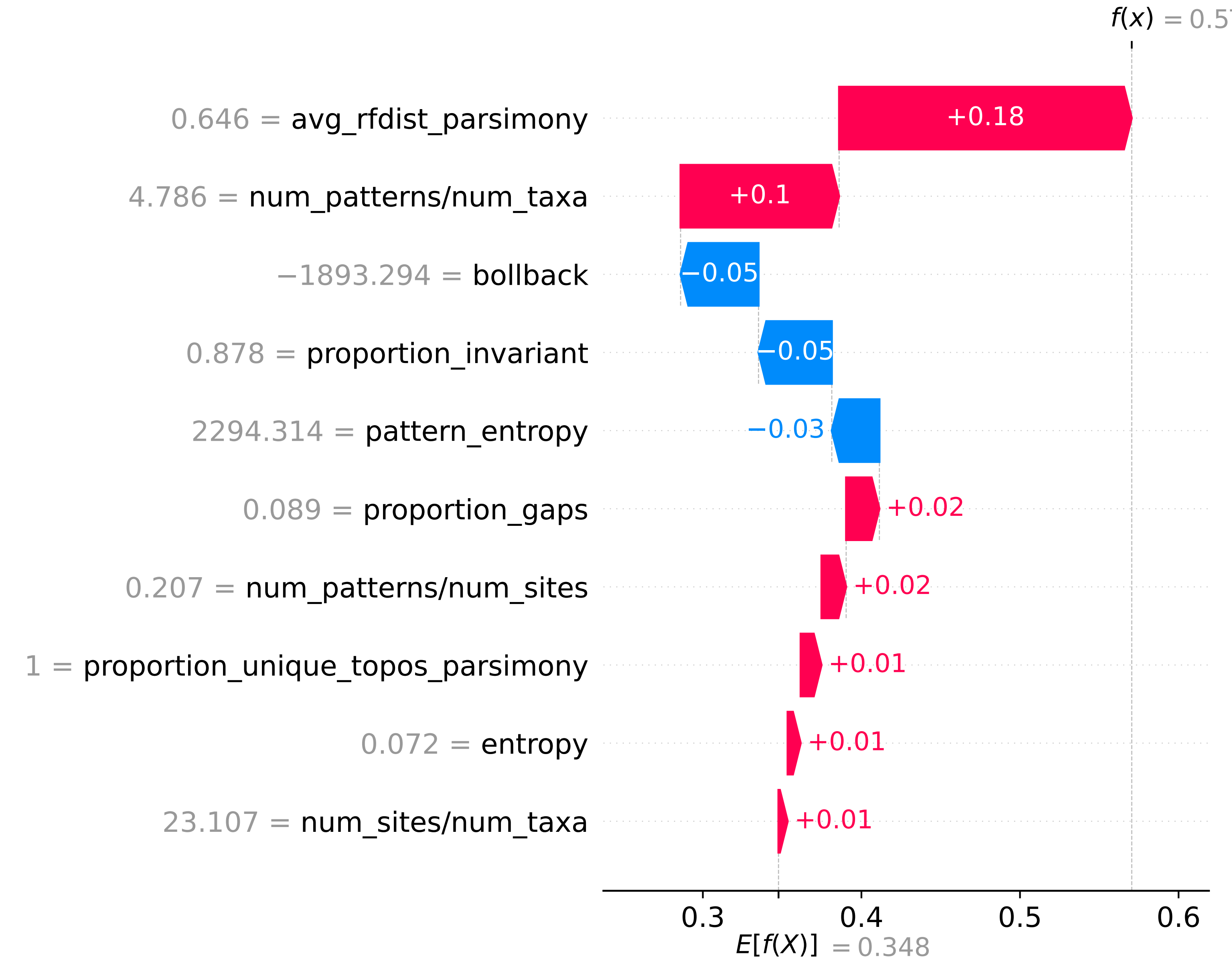

### SupplementaryFigure_5.pdf

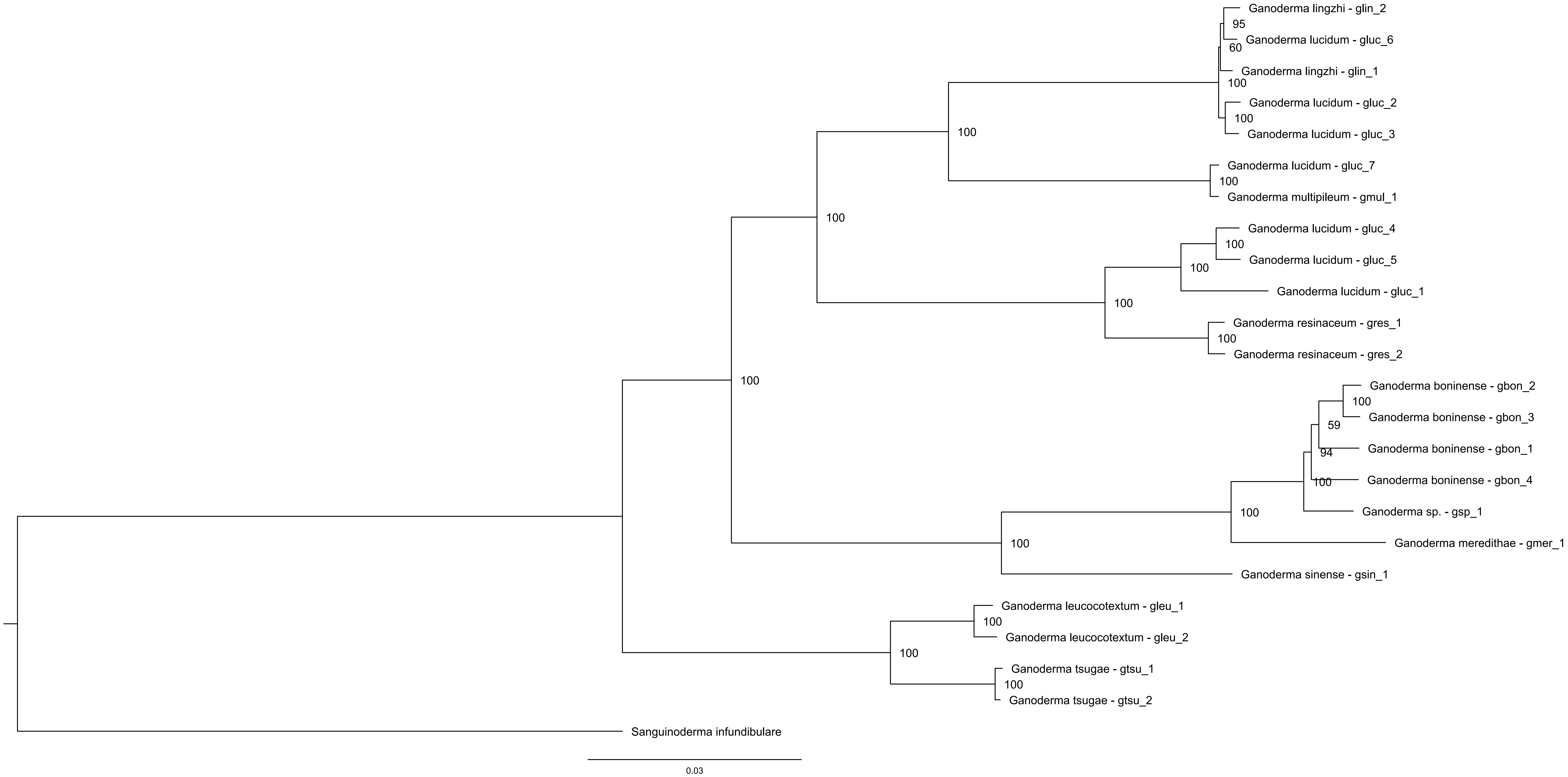
